# Brief sensory deprivation triggers cell type-specific structural and functional plasticity in olfactory bulb neurons

**DOI:** 10.1101/2020.05.10.086926

**Authors:** Elisa Galliano, Christiane Hahn, Lorcan Browne, Paula R. Villamayor, Matthew S. Grubb

## Abstract

Can alterations in experience trigger different plastic modifications in neuronal structure and function, and if so, how do they integrate at the cellular level? To address this question, we interrogated circuitry in the mouse olfactory bulb responsible for the earliest steps in odour processing. We induced experience-dependent plasticity in mice by blocking one nostril for a day, a minimally-invasive manipulation which leaves the sensory organ undamaged and is akin to the natural transient blockage suffered during common mild rhinal infections. We found that such brief sensory deprivation produced structural and functional plasticity in one highly specialised bulbar cell type: axon-bearing dopaminergic neurons in the glomerular layer. After 24h naris occlusion, the axon initial segment (AIS) in bulbar dopaminergic neurons became significantly shorter, a structural modification that was also associated with a decrease in intrinsic excitability. These effects were specific to the AIS-positive dopaminergic subpopulation, because no experience-dependent alterations in intrinsic excitability were observed in AIS-negative dopaminergic cells. Moreover, 24h naris occlusion produced no structural changes at the AIS of bulbar excitatory neurons – mitral/tufted and external tufted cells – nor did it alter their intrinsic excitability. By targeting excitability in one specialised dopaminergic subpopulation, experience-dependent plasticity in early olfactory networks might act to fine-tune sensory processing in the face of continually fluctuating inputs.

## INTRODUCTION

One way that animals can ensure appropriate behavioural choices when faced with an ever-changing environment is to alter the way they process sensory inputs. To implement such adaptive control at the level of neuronal networks, a huge range of cellular mechanisms of neuronal plasticity have been elucidated. These include structural changes in neuronal morphology, functional changes of synaptic strength, and/or modulation of the machinery that sustains a neuron’s intrinsic excitability (Brzosko et al., 2019; Citri and Malenka, 2008; Debanne et al., 2019; Kullmann et al., 2012; Wefelmeyer et al., 2016). This extensive repertoire also includes a form of structural plasticity tightly linked with changes in neuronal excitability: plasticity of the axon initial segment (AIS).

Structurally, the AIS is a subcellular zone located in the proximal portion of the axon, where an intricate arrangement of cytoskeletal and scaffolding proteins anchors a membrane-bound collection of signaling molecules, receptors and ion channels (Hamdan et al., 2020; Leterrier, 2018; Vassilopoulos et al., 2019). Functionally, the AIS serves two key roles: maintenance of dendritic/axonal polarity (Hedstrom et al., 2008), and initiation of action potentials (Bean, 2007; Kole et al., 2007). Plastically, the AIS has been proven capable of changing its structure in terms of length, distance from the soma, and/or molecular content (Ding et al., 2018; Grubb and Burrone, 2010; Kuba et al., 2010, 2015; Lezmy et al., 2017).

How is AIS plasticity driven by changes in neuronal activity? *In vitro*, elevated activity causes the AIS of excitatory neurons to relocate distally or to decrease in length, structural changes associated with decreased functional excitability (Evans et al., 2013, 2015; Grubb and Burrone, 2010; Horschitz et al., 2015; Lezmy et al., 2017; Muir and Kittler, 2014; Sohn et al., 2019; Wefelmeyer et al., 2015). *In vivo*, various groups have described activity-dependent structural AIS plasticity in excitatory neurons, usually induced by manipulations that are long in duration and/or involve damage to peripheral sensory organs ((Gutzmann et al., 2014; Kuba et al., 2010; Pan-Vazquez et al., 2020), but see (Jamann et al., 2020)). But is AIS plasticity a prerogative of excitatory neurons, or is it also included in the plasticity toolkit of inhibitory cells? We previously found that, *in vitro*, inhibitory dopaminergic (DA) interneurons in the olfactory bulb (OB) are capable of bidirectional AIS plasticity, inverted in sign with respect to their excitatory counterparts: their AIS increases in length and relocates proximally in response to chronic depolarization, and shortens when spontaneous activity is silenced (Chand et al., 2015). Taken together, these studies begin to paint a picture of how different cell types respond to changes in incoming activity levels by initiating distinct plastic structural changes at their AIS. However, many key questions remain unanswered. Are more physiological, minimally-invasive sensory manipulations sufficient to induce AIS plasticity *in vivo?* In the intact animal, can AIS plasticity occur over more rapid timescales? And do excitatory and inhibitory neurons in sensory circuits respond to such brief and naturally-relevant sensory manipulation with similar levels of AIS plasticity?

To address these questions, we interrogated circuitry in the mouse OB responsible for the earliest steps in odour processing (Shepherd, 2005). At just one synapse away from the sensory periphery, activity in the OB can be readily and reliably altered by physiologically-relevant alterations in sensory experience (Coppola, 2012). In our case this was achieved by unilaterally plugging a nostril for just one day, a minimally-invasive manipulation which effectively mimics the sensory disturbance associated with common respiratory infections without damaging the olfactory sensory epithelium (Fokkens et al., 2012). We found that such brief sensory deprivation produced structural and functional plasticity only in axon-bearing dopaminergic (DA) neurons in the bulb’s glomerular layer (Chand et al., 2015; Galliano et al., 2018). By targeting excitability in one specialised dopaminergic subpopulation, experience-dependent plasticity in early olfactory networks might act to fine-tune sensory processing in the face of continually fluctuating inputs.

## MATERIALS AND METHODS

### Animals

We used mice of either gender, and housed them under a 12-h light-dark cycle in an environmentally controlled room with free access to water and food. Wild-type C57Bl6 mice (Charles River) were used either as experimental animals, or to back-cross each generation of transgenic animals. The founders of our transgenic mouse lines – DAT-Cre (B6.SJL-Slc6a3^*tm1.1(cre)Bkmn*/J^, Jax stock 006660) and flex-tdTomato (B6.Cg–Gt(ROSA)^26Sortm9(CAG-tdTomato)Hze^, Jax stock 007909) were purchased from Jackson Laboratories. All experiments were performed between postnatal days (P) 21 and 35. All experiments were performed at King’s College London under the auspices of UK Home Office personal and project licences held by the authors.

### Sensory manipulation

To perform unilateral naris occlusion, mice were briefly anaesthetised (<5 min) with isoflurane. In the occluded group, a custom-made ^~^5 mm Vaseline-lubricated plug, constructed by knotting suture (Ethylon polymide size 6, non-absorbable suture, Ethicon, UK) around a piece of unscented dental floss and pulled through the lumen of PTFE-tubing with an outer-diameter of 0.3 mm and inner-diameter of 0.3 mm (VWR International, cat#: S1810-04; see (Cummings et al., 2014)) was inserted into the right nostril and left for 24 hours. Only the right olfactory bulb was then used for experiments. At the termination of each experiment, post-hoc visual observation of the nasal cavity was always performed to ensure that the plug had remained in place. The few mice where the plug could not be found were not used for experiments. All control animals were gender and age-matched mice left unperturbed in their home cage. For both control and occluded groups, only right bulbs were analysed.

### Immunohistochemistry

Mice were anesthetized with an overdose of pentobarbital and then perfused with 20 mL PBS with heparin (20 units.mL^−1^), followed by 20mL of 1% paraformaldehyde (PFA; TAAB Laboratories; in 3% sucrose, 60 mM PIPES, 25 mM HEPES, 5 mM EGTA, and 1 mM MgCl_2_).

To expose the olfactory epithelia the rostral half of the calvaria (anterior to the bregma) and the nasal bone were removed, and the samples were first post-fixed overnight (4°C) and then placed in 0.25 M EDTA (Invitrogen AM9261) in PBS at 4°C for 3 days for decalcification. After overnight cryoprotective treatment with 30% sucrose (Sigma S9378), they were then embedded in OCT (VWR Chemicals 00411243), frozen in liquid nitrogen and sliced on a cryostat (Leica CM 1950) into 20 μm slices.

The olfactory bulbs were dissected and post-fixed in 1% PFA for 2-7 days, then embedded in 5% agarose and sliced at 50 μm using a vibratome (VT1000S, Leica). For experiments which aimed at comparing intensity of staining across mice, we co-embedded the bulbs of one control and one occluded mouse in a large agarose block (“set”), and from then forward we processed them as a unit (Vlug et al., 2005). Free-floating slices or sets were washed with PBS and incubated in 5% normal goat serum (NGS) in PBS/Triton/azide (0.25% triton, 0.02% azide) for 2 h at room temperature. They were then incubated in primary antibody solution (in PBS/Triton/azide) for 2 days at 4°C. The primary antibodies used and their respective concentrations are indicated in the table below.

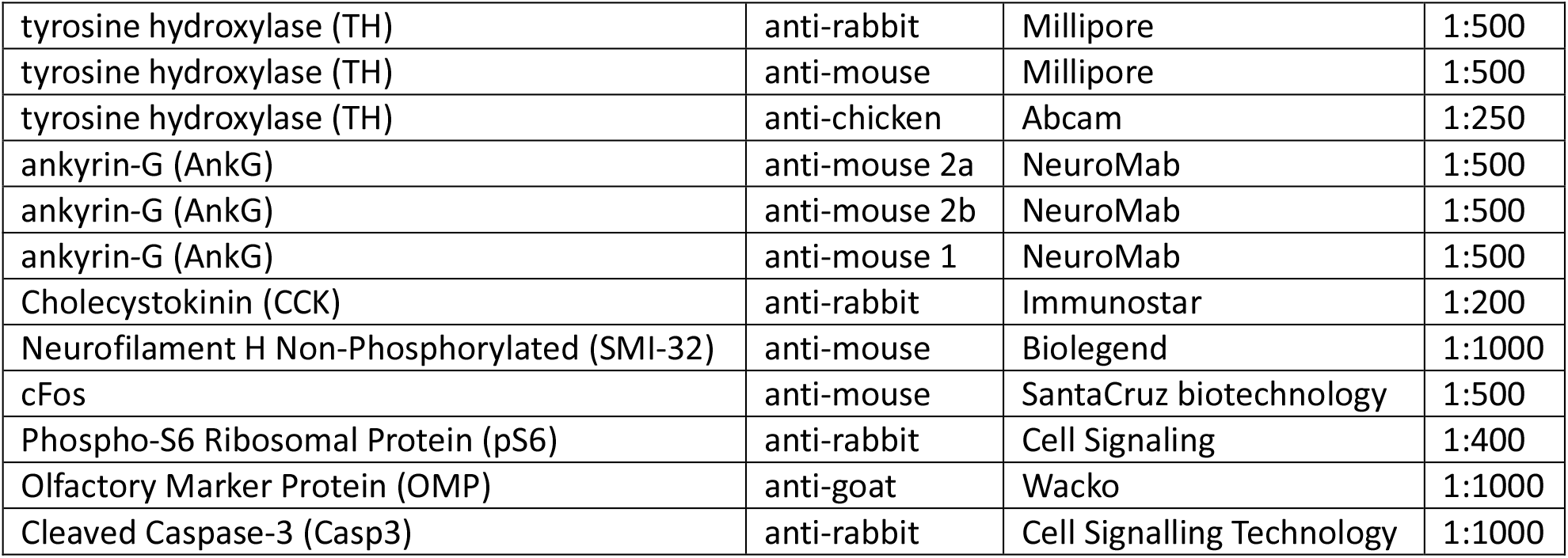

Slices were then washed three times for 5 min with PBS, before being incubated in secondary antibody solution (species-appropriate Life Technologies Alexa Fluor^®^; 1:1000 in PBS/Triton/azide) for 3 h at room temperature. After washing in PBS, slices were either directly mounted on glass slides, Menzel-Gläser) with MOWIOL-488 (Calbiochem), or first underwent additional counterstaining steps with NucRed Live 647 (Invitrogen R37106) at room temperature for 25 min to visualize cell nuclei, or with 0.2% sudan black in 70% ethanol at room temperature for 3 min to minimize autofluorescence. Unless stated otherwise all reagents were purchased from Sigma.

### Fixed-tissue imaging and analysis

All images were acquired with a laser scanning confocal microscope (Zeiss LSM 710) using appropriate excitation and emission filters, a pinhole of 1 AU and a 40x oil immersion objective. Laser power and gain were set to either prevent signal saturation in channels imaged for localisation analyses, or to permit clear delineation of neuronal processes in channels imaged for neurite identification (*e.g.*, TH, SMI-32, CCK). All quantitative analysis was performed with Fiji (Image J) by experimenters blind to group identity.

For olfactory epithelium (OE) analysis, 4 images were acquired from consistently positioned septal and dorsomedial regions of interest within each section, with a 1x zoom (0.415 μm/pixel), 512×512 pixels, and in z-stacks with 1 μm steps. OE thickness was measured on single plane images by drawing a straight line, parallel to olfactory sensory neurons (OSNs) dendrites, from the lamina propria to the tips of the OSN dendrites (visualized with OMP label). OSN density was calculated on single-plane image by counting the number of clearly OMP-positive somas (OMP label surrounding NucRed+ nucleus), divided by length of OE in that image, x100 for comparative purposes (Cheetham et al., 2016; Kikuta et al., 2015). To quantify cell apoptosis, expressed as cells/mm for comparative purposes, the number of Caspase-3 positive cells was measured from Z-stacks through entire slice, and then divided by total length of the OE in the stack (OE length x n of Z steps) (Kikuta et al., 2015).

For activity early genes and TH expression in the olfactory bulb, images were taken with a 1x zoom (0.415 μm/pixel), 512×512 pixels, and in z-stacks with 1 μm steps, with identical laser power and digital gain/offset settings within each set. To avoid selection biases, all cells present in the stack and positive for the identifying marker (TH or SMI-32) were measured. DA cell density was calculated for each image by dividing the number of analysed TH-positive cells by the volume of the glomerular layer (z depth x glomerular layer area, drawn and measured in a maximum intensity projection of the TH channel). SMI-32 positive M/TCs were selected by position in the mitral layer; SMI-32 positive ETCs were included in the analysis only if their soma bordered with both the GL and external plexiform layer. TH positive DA cells were included in the analysis only if their soma was in or bordering with the glomerular layer. Soma area was measured at the single plane including the cell’s maximum diameter by drawing a region of interest (ROI) with the free-hand drawing tool. Within each co-embedded set, the staining intensity of each ROI (expressed as mean grey value) was normalized to the mean value of staining intensity across all measured cells in the control slice. Staining intensity in AIS-positive DA cells (*i.e.*, AnkG+/TH+) was normalized to the average TH or cFos staining of the overall DA cell population in the set control slice.

For AIS identification, images were taken with 3x zoom, 512×512 pixels (0.138 μm/pixel) and in z-stacks with 0.45 μm steps. While in all glutamatergic neurons only one extensive AnkG-positive region could be found on the proximal part of a process originating directly from the soma, DA cells’ AISes were found either on processes originating directly from the soma (“soma-origin”) or on a process that did not originate directly from the soma (“dendrite-origin”). Moreover, as previously reported in the literature (Kosaka et al., 2008; Meyer and Wahle, 1988) a minority of DA cells was found to carry multiple AISes (10% of all imaged cells) and excluded from further analysis. In all cells carrying a single AIS, its distance from soma and length were measured in Fiji/ImageJ using the View5D plugin, which allows for 3D manual tracing of cell processes. Laser power and gain settings were adjusted to prevent signal saturation in the AIS label AnkG; cellular marker TH or SMI-32 signal was usually saturated to enable clear delineation of the axon. The AIS distance from soma was traced from the start of the axon (identified as the tapering of the soma into the axon hillock) to the point where AnkG staining became clearly identifiable. The AIS length was similarly traced by following the AnkG staining along the axon course. To confirm the reliability of this manual tracing method, a subset of 50 AISes was analyzed twice by EG, blindly and with two weeks’ inter-analysis interval. Measurements of both distance from soma and length were highly consistent between the two analysis sessions (AIS distance from soma: difference 0.006 ± 0.097 μm, r^2^=0.75; AIS length: 0.139 ± 0.195 μm, r^2^=0.95).

### Acute-slice electrophysiology

P21-35 C57Bl6 or DAT^*Cre*^-tdTomato mice were decapitated under isoflurane anaesthesia and the OB was removed and transferred into ice-cold slicing medium containing (in mM): 240 sucrose, 5 KCl, 1.25 Na_2_HPO_4_, 2 MgSO_4_, 1 CaCl_2_, 26 NaHCO_3_ and 10 D-Glucose, bubbled with 95% O_2_ and 5% CO_2_. Horizontal slices (300 μm thick) of the olfactory bulb were cut using a vibratome (VT1000S, Leica) and maintained in ACSF containing (in mM): 124 NaCl, 5 KCl, 1.25 Na_2_HPO_4_, 2 MgSO_4_, 2 CaCl_2_, 26 NaHCO_3_ and 20 D-Glucose, bubbled with 95% O_2_ and 5% CO_2_ for >1 h before experiments began.

Whole-cell patch-clamp recordings were performed using an Axopatch amplifier 700B (Molecular Devices, Union City, CA, USA) at physiologically-relevant temperature (32-34°C) with an in-line heater (TC-344B, Warner Instruments). Signals were digitized (Digidata 1550, Molecular Devices) and Bessel-filtered at 3 KHz. Recordings were excluded if series (RS) or input (RI) resistances (assessed by −10 mV voltage steps following each test pulse, acquisition rate 20KHz) were respectively bigger than 30 MΩ or smaller than 100 MΩ for DA neurons, bigger than 30 MΩ or smaller than 30 MΩ for ETCs, bigger than 20 MΩ or smaller than 40 MΩ for M/TCs, or if they varied by > 20% over the course of the experiment. Fast capacitance was compensated in the on-cell configuration and slow capacitance was compensated after rupture. Cell capacitance (Cm) was calculated by measuring the area under the curve of the transient capacitive current elicited by a −10 mV voltage step. Resting membrane potential (Vm) was assessed immediately after break-in by reading the voltage value in the absence of current injection (I=0 configuration). Recording electrodes (GT100T-10, Harvard Apparatus) were pulled with a vertical puller (PC-10, Narishige) and filled with an intracellular solution containing (in mM): 124 K-Gluconate, 9 KCl, 10 KOH, 4 NaCl, 10 HEPES, 28.5 Sucrose, 4 Na_2_ATP, 0.4 Na_3_GTP (pH 7.25-7.35; 290 MOsm) and Alexa 488 (1:150). Cells were visualized using an upright microscope (Axioskop Eclipse FN1 Nikon, Tokyo, Japan) equipped with a 40X water immersion objective, and for DA cell identification tdT fluorescence was revealed by LED (CoolLED pE-100) excitation with appropriate excitation and emission filters (ET575/50m, CAIRN Research, UK). M/TCs were identified based on location in the mitral layer and large somas. ETCs were identified based on: (a) location in the lower glomerular layer / upper external plexiform layer; (b) large and balloon-shaped soma and, often, visible large apical dendrite; (c) characteristic spontaneous burst firing when unclamped; (d) an relatively depolarized resting membrane potential of ^~^−55 mV; and (e) distinct depolarising sag potential when injected with prolonged negative current steps in current clamp mode (Liu and Shipley, 2008; Liu et al., 2013).

In current-clamp mode, evoked spikes were measured with *V*_hold_ set to −60 ± 3 mV for M/TCs and DA cells, and to −55 ± 3 mV for ETCs. For action potential waveform measures, we injected 10-ms-duration current steps from 0 pA of increasing amplitude (Δ5/20 pA) until we reached the current threshold at which the neuron reliably fired an action potential (*V*_m_ > 0 mV; acquisition rate 200 KHz). For multiple spiking measures, we injected 500-ms-duration current steps from 0pA of increasing amplitude (Δ2/10 pA) until the neuron passed its maximum firing frequency (acquisition rate 50 KHz). Exported traces were analyzed using either ClampFit (pClam10, Molecular Devices) or custom-written routines in MATLAB (Mathworks). Before differentiation for *dV/dt* and associated phase plane plot analyses, recordings at high temporal resolution (5 μs sample interval) were smoothed using a 20 point (100 μs) sliding filter. Voltage threshold was taken as the potential at which *dV/dt* first passed 10 V/s. Onset rapidness was taken from the slope of a linear fit to the phase plane plot at voltage threshold. For DA cells, monophasic versus biphasic phase plane plots were visually determined by EG and MSG. We classified completely monotonic plots with continually increasing rate-of-rise as monophasic, and any plots showing a clear inflection in rate-of-rise over the initial rising phase as biphasic. Any discrepancies in classification were resolved by mutual agreement. We also corroborated our subjective classification using a quantitative measure of spike onset sharpness: the ratio of errors produced by linear and exponential fits to the peri-threshold portion of the phase plane plot (Baranauskas et al., 2010; Volgushev et al., 2008). Spike width was measured at the midpoint between voltage threshold and maximum voltage. Rheobase and afterhyperpolarization values were both measured from responses to 500 ms current injection, the latter from the local voltage minimum after the first spike fired at rheobase. Input-output curves were constructed by simply counting the number of spikes fired at each level of injected current.

### Statistical analysis

Statistical analysis was carried out using Prism (Graphpad), SPSS (IBM) or Matlab (Mathworks). Sample distributions were assessed for normality with the D’Agostino and Pearson omnibus test, and parametric or non-parametric tests carried out accordingly. α values were set to 0.05, and all comparisons were two-tailed. For multilevel analyses, non-normal distributions were rendered normal by logarithmic transform. These parameters were then analysed using linear mixed models (SPSS) with mouse or set as the subject variable (Aarts et al., 2014).

## RESULTS

### Brief unilateral naris occlusion leaves the olfactory epithelium undamaged

Olfactory sensory deprivation in mice can be achieved surgically by cauterisation of one naris, or mechanically by insertion of a custom-made and removable nasal plug (Coppola, 2012). Traditionally, both methods have been employed for prolonged periods (weeks, months at a time), and are accompanied by pronounced and widespread changes in olfactory bulb architecture, including overall OB size. This scenario is potentially pathological, and does not reflect the most common deprivation that this sensory system has to deal with: a nasal blockage lasting less than 5 days (Fokkens et al., 2012).

In order to induce activity-dependent plasticity within a more naturally-relevant timeframe, we therefore employed the custom-made plug method, but left the plug in place for just one day (Fig. 1A)(Cummings and Brunjes, 1997). Because of concerns regarding abnormal airflow through the remaining open nostril in unilaterally occluded animals (Coppola, 2012; Kass et al., 2013; Wu et al., 2017), we did not compare open and occluded hemispheres within the same experimental animals. Instead, juvenile (P27) wild-type mice were either left unperturbed (Fig. 1A; control group, Ctrl, black) or had one nostril plugged for 24 h (occluded group, Occl, orange), before being perfused and processed for immunohistochemistry.

**Figure 1.**
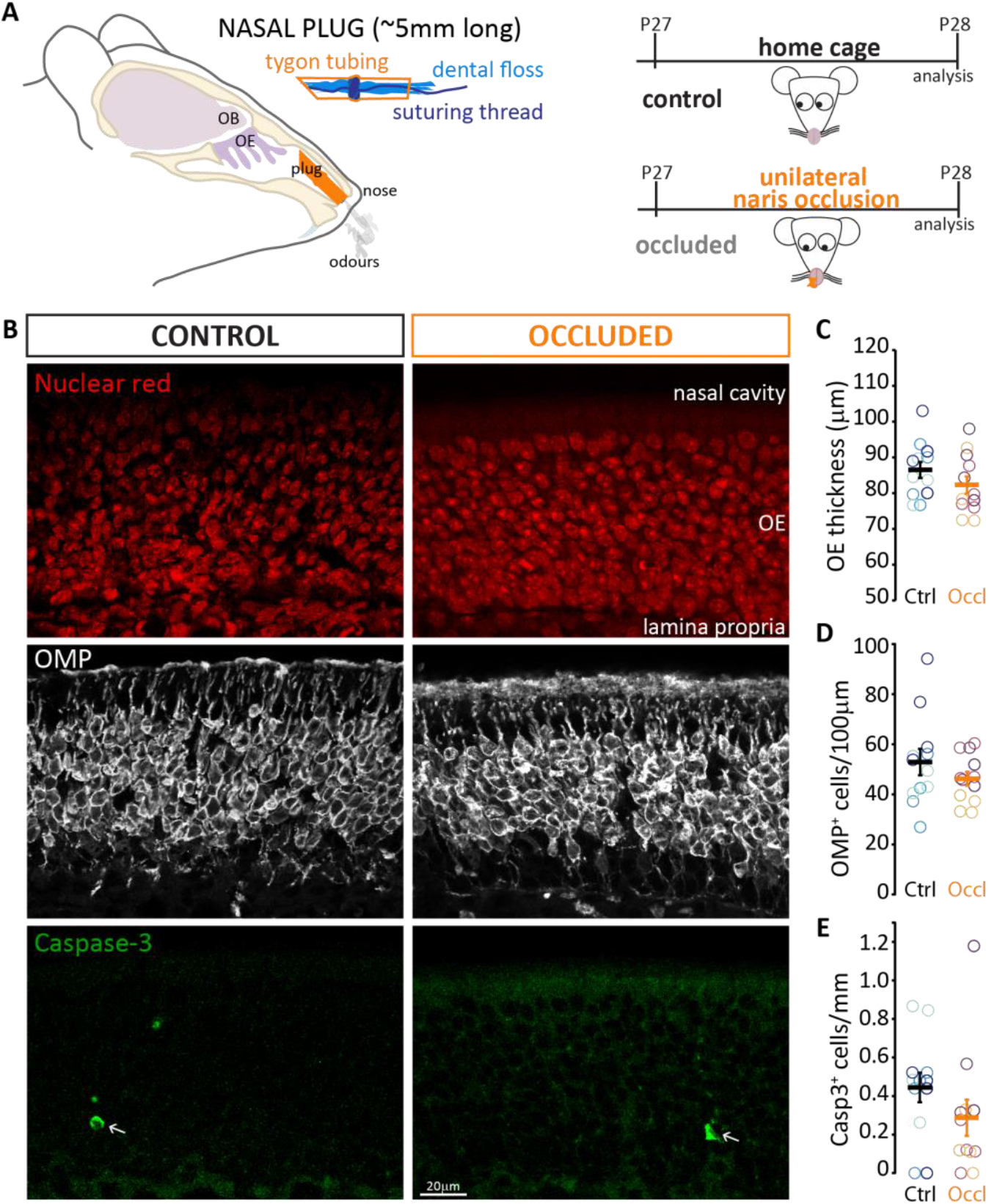
Brief unilateral naris occlusion does not damage the olfactory epithelium. **(A)** Left: schematic representation of the custom-made plug (orange), and of how it lodges in the mouse nasal cavity blocking air flow without direct contact with the olfactory epithelium (OE). OB, olfactory bulb. Right: timeline of sensory manipulation. P27 littermates were either left undisturbed in their home cage (control group, black), or had one naris plugged for 24h (occluded group, orange). All mice were then processed for immunohistochemistry at P28. **(B)** Example images of olfactory epithelia in control and occluded mice. Red: nuclear marker; grey: OMP, olfactory marker protein; green: activated Caspase-3, apoptosis marker. Arrow indicates rare caspase-positive cells. **(C)** Thickness of the olfactory epithelium in control (Ctrl, black, mean+SEM 86.51 ± 2.26 μm) and occluded (Occl, orange, mean+SEM 82.26 + 2.40 μm) mice, p=0.19. **(D)** Density of OMP-positive cells in control (52.84 ± 5.24 cells/100 μm) and occluded (46.20 + 2.78 cells/100 μm) mice, p=0.47. **(E)** Density of Caspase-3-positive cells in control (0.39 ± 0.084 cells/mm) and occluded (0.29 ± 0.094 cells/mm) mice, p=0.54. Empty circles represent individual sample regions; different colours indicate different mice (n=12, N=3 for both Ctrl and Occl); thick line shows mean ± SEM.

To confirm the expected lack of peripheral pathology with this approach (Cheetham et al., 2016; Kikuta et al., 2015), we assessed the impact of plug insertion on the olfactory epithelium (OE; Fig. 1B). We found no difference between control and 24 h-occluded groups in overall OE thickness (Fig. 1C; Ctrl mean ± SEM 86.51 ± 2.26 μm; Occl 82.26 ± 2.40 μm; mixed model ANOVA nested on mouse, effect of treatment F_1,24_ = 1.81, p = 0.19). Similarly, the density of olfactory sensory neurons (OSNs, identified by immunolabel for olfactory marker protein, OMP) did not differ between control and occluded mice (Fig. 1D; Ctrl mean ± SEM 52.84 ± 5.24 cells/100 μm; Occl 46.20 ± 2.78 cells/100 μm; mixed model ANOVA nested on mouse, effect of treatment F_1,6_ = .584, p = 0.47), nor did the density of apoptotic cells positive for activated Caspase-3 (Fig. 1E; Ctrl mean ± SEM 0.39 ± 0.084 cells/mm; Occl 0.29 ± 0.094 cells/mm; mixed model ANOVA nested on mouse, effect of treatment F_1,6_ = .423, p = 0.54). Overall, these data suggest that brief olfactory deprivation carried out with a custom-made plug has no impact on the overall structure and health of the olfactory epithelium.

### Brief unilateral naris occlusion alters the activity of inhibitory and excitatory bulbar neurons

Given that our chosen sensory manipulation is well within naturally-experienced timeframes (Fokkens et al., 2012) and does not overtly damage the peripheral sense organ, we next checked that it was effective in reducing ongoing activity levels in downstream OB neurons.

We processed the OBs of control and occluded mice to quantify the expression of activity markers with immunohistochemistry. To control for differences in antibody exposure, we co-embedded slices from control and occluded mice in agarose blocks (“sets”, Fig. 2A) for consistent histological processing, and normalized activity marker intensity within each set (see Methods).

**Figure 2.**
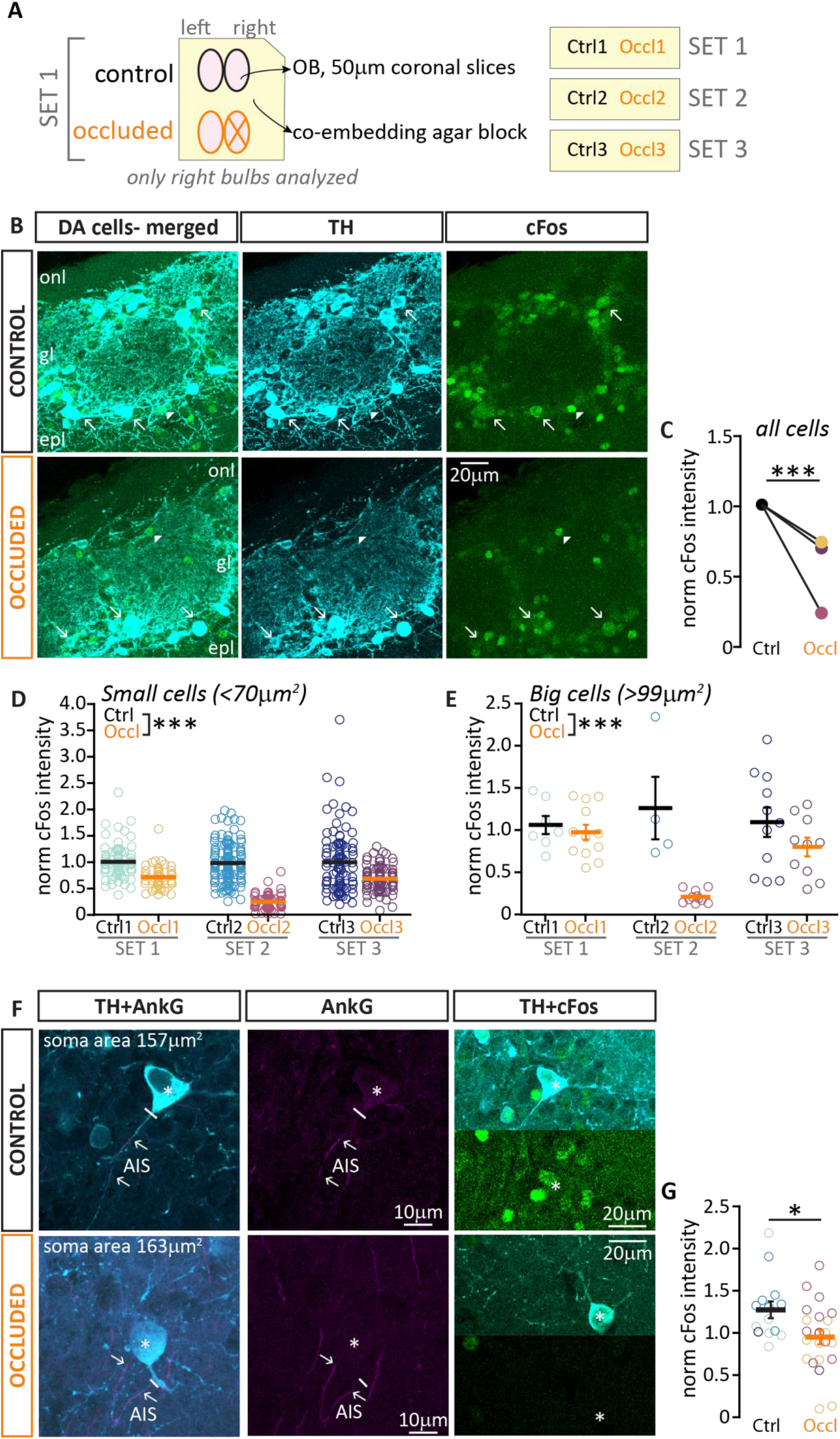
Brief unilateral naris occlusion decreases activity levels in both major subtypes of olfactory bulb dopaminergic neurons. **(A)** Schematic representation of the immunohistochemistry procedure: 50 μm coronal slices of OBs from one control and one occluded (X) mouse were co-embedded in an agarose block (“set”) and processed together. Each set was analysed independently and staining intensity was normalized to the average of control measurements – see Methods. **(B)** Example maximum intensity projection image of dopaminergic neurons (DA cells, cyan) visualized via anti-tyrosine hydroxylase (TH) staining, and label for the activity early gene cFos (green), in control (black) and occluded (orange) mice, onl = olfactory nerve layer; gl = glomerular layer; epl = external plexiform layer. Arrows indicate cFos positive DA cells, arrowheads indicate TH-negative/cFos-positive cells. **(C)** Average normalised cFos intensity in TH-positive cells (of any soma size) in control (n=369; mean±SEM 1 ± 0.02) and occluded (n=301, 0.56 ± 0.02) mice. **(D)** Normalized cFos intensity in TH-positive cells with soma area < 70 μm^2^ (putative anaxonic DA cells), from 3 sets of Ctrl (black, n=298, mean±SEM 0.99 ± 0.03) and Occl (orange, n=192, 0.56 ± 0.02) mice. **(E)** Normalized cFos intensity in TH-positive cells with soma size > 99 μm^2^ (putative axonal DA cells), from 3 sets of Ctrl (black, n=22, mean±SEM 1.11 ± 0.11) and Occl (orange, n=33, 0.67 ± 0.07) mice. **(F)** Example images of cFos expression (green) in DA cells (TH-positive; cyan) with an identified ankyrin-G (AnkG; magenta)-positive AIS (arrows). The solid line indicates the emergence of the axonal process from the soma (asterisk). **(G)** Normalized cFos intensity in AnkG-positive/TH-positive cells in control (n=14, mean±SEM 1.28 ± 0.10) and occluded (n=22, 95.18 ± 0.09) mice. Empty circles represent individual cells and different colours indicate different mice (N=3 for both Ctrl and Occl); thick lines show mean ± SEM; *= p<0.05, ***= p<0.0001.

We first analyzed the expression of the immediate early gene cFos (Barnes et al., 2015) in dopaminergic inhibitory neurons (DA cells, identified via tyrosine hydroxylase, TH, immunoreactivity; Fig. 2B). DA cells in occluded bulbs displayed markedly and consistently lower cFos levels than their co-embedded control counterparts, and this effect was highly significant in multilevel statistical analyses that account for inter-set variation (Fig. 3B; Ctrl mean ± SEM 1 ± 0.02; Occl 0.56 ± 0.02; mixed model ANOVA nested on set, p<0.0001).

**Figure 3.**
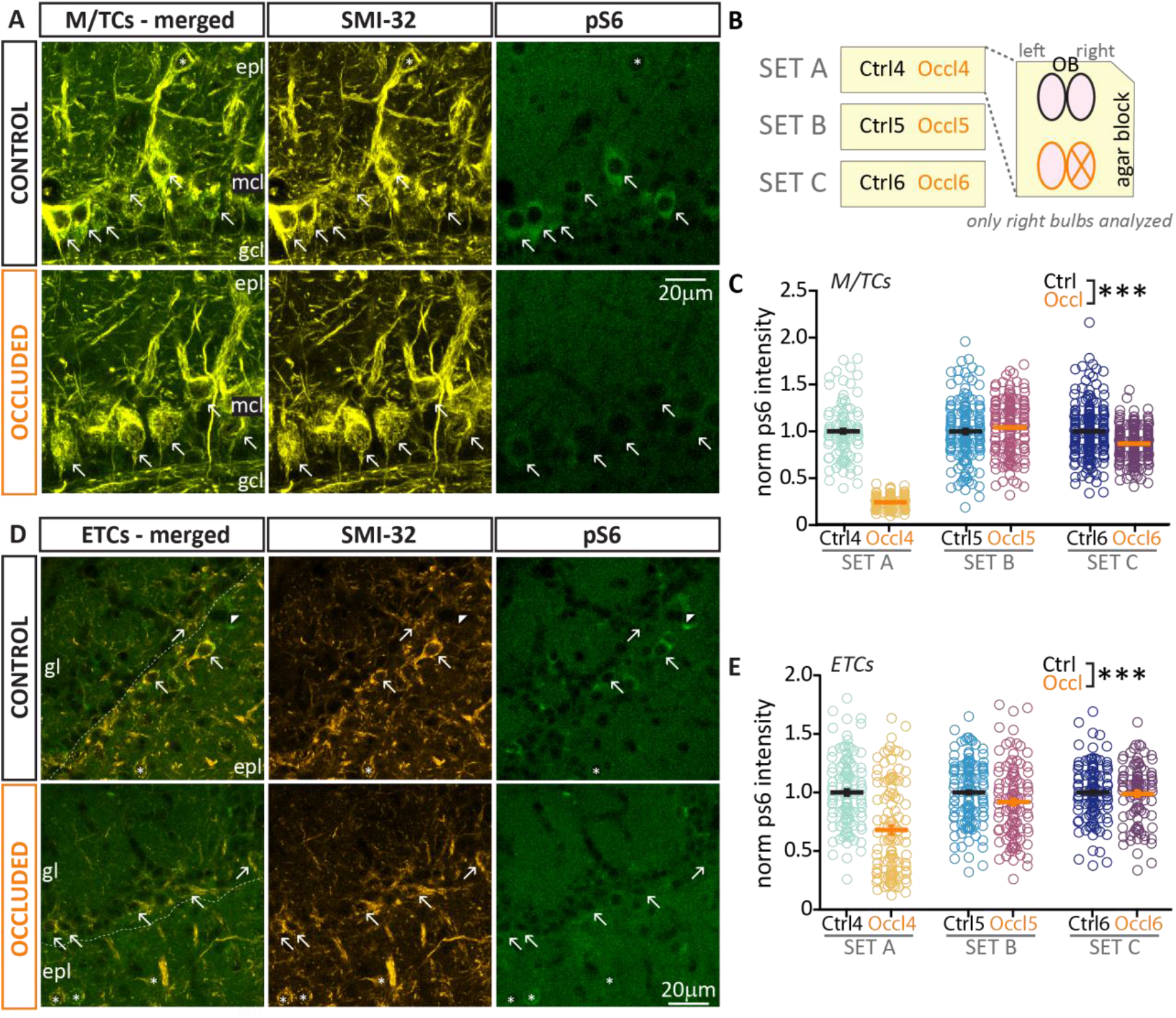
Brief unilateral naris occlusion decreases activity levels in bulbar excitatory neurons. **(A)** Example maximum intensity projection image of bulbar mitral/tufted cells (M/TCs, visualized via SMI-32 staining, yellow) and activity early gene pS6 (green), epl = external plexiform layer; mcl = mitral cell layer; gcl = granule cells layer. **(B)** Schematic representation of the immunohistochemistry procedure: 50 μm coronal slices of OBs from one control and one occluded (X) mouse were co-embedded in an agarose block (“set”) and processed together. Each set was analysed independently and staining intensity was normalized to the average of control measurements – see Methods. **(C)** Normalized pS6 intensity in M/TCs from 3 sets of Ctrl (black, n=451, mean+SEM 1 ± 0.01) and Occl (orange, n=478, 0.76 + 0.02) mice, p<0.0001). Arrows indicate pS6 positive cells. **(D-E)** As in A and C, example maximum intensity projection image and analysis of normalised pS6 intensity in external tufted cells (ETCs, amber) from 3 sets of Ctrl (black, n=362, mean±SEM 1 ± 0.01) and Occl (orange, n=371, 0.87 ± 0.02) mice; p<0.0001. Only SMI-32 positive cells located in the glomerular layer (gl) were included in the analysis (arrows), while SMI-32 positive cells located in the epl (asterisks) were excluded. Empty circles represent individual cells and different colours indicate different mice (N=3 for both Ctrl and Occl); thick line shows mean ± SEM.

Previous work from ourselves and others has found that bulbar DA neurons are a heterogeneous population (Chand et al., 2015; Galliano et al., 2018; Kosaka et al., 2019). Two non-overlapping subtypes can be identified by a spectrum of different morphological and functional characteristics, as well as by a binary classifier: the presence or absence of an axon and its key component, the axon initial segment (AIS) (Chand et al., 2015; Galliano et al., 2018). So, does brief unilateral naris occlusion downregulate activity in both axon-bearing and anaxonic DA subtypes? Soma size is a readily-obtainable proxy indicator for DA subtypes: anaxonic DA cells are usually small, while axon-bearing DA cells tend to have very large somas. Using previously defined lower (<70 μm^2^) and upper (>99μm^2^) bounds of the OB DA soma size distribution (Galliano et al., 2018), we found that both small/putative anaxonic DA cells and large/putative axon-bearing DA cells from occluded mice display reduced cFos staining relative to their co-embedded control counterparts. Although the smaller sample size of the much rarer large DA cells accentuated variability across staining sets here, this effect was highly significant for both cell types in analyses that specifically account for that variation (small cells: Ctrl mean ± SEM 0.99 ± 0.03, Occl 0.56 ± 0.02; big cells: Ctrl mean ± SEM 1.11 ± 0.11, Occl 0.67 ± 0.07; for both, mixed model ANOVA nested on mouse, p<0.0001). Finally, to further confirm these results in DA cells which definitively belonged to the axon-bearing subtype, we co-stained a subset of tissue with the AIS marker ankyrin-G (AnkG) and measured cFos levels in AnkG+/TH+ DA cells (Fig. 3E; see Methods). Once more, we found significantly dimmer cFos fluorescence in occluded cells (Fig.3F; Ctrl mean ± SEM 1.28 ± 0.10, Occl 95.18 ± 0.09, Mann-Whitney, p=0.02).

This effect of naris occlusion on activity levels was more variable, but nevertheless also present overall in bulbar glutamatergic neurons. These belong to two main classes defined by location and axonal projections: mitral/tufted cells and external tufted cells. Mitral/tufted cells (M/TCs, Fig. 3A), whose soma sits in the mitral cell layer, are the bulbar network’s principal neurons; they extend their apical dendrites to the glomerular layer where they receive direct and indirect inputs from OSNs, and send their axons to higher olfactory areas, including piriform cortex (Imai, 2014). External tufted cells (ETCs, Fig. 3C) are glutamatergic interneurons located in the glomerular layer, where they provide local dendrodendritic amplification of sensory inputs (Gire et al., 2012; Najac et al., 2011). ETC axons do not leave the OB, but target deep-layer networks beneath sister glomeruli in the opposite hemi-bulb (Cummings and Belluscio, 2010; Lodovichi et al., 2003). To identify both classes of excitatory neurons, we labelled bulbar slices with the neurofilament marker protein H, clone SMI-32 (see Methods).

We co-stained with antibodies against another activity marker, phospho-S6 ribosomal protein (pS6; Knight et al., 2012) which in bulbar glutamatergic cells gives higher intensity and consistency of staining than cFos (Fig. 3A-C). We found that both in M/TCs and ETC from occluded slices, the relative intensity levels of pS6 were markedly variable across staining sets. Nevertheless, multilevel analyses that account for this variability found that pS6 intensity was significantly decreased overall in both cell types in occluded bulbs when compared to co-embedded controls (M/TC Occl mean ± SEM 0.76 ± 0.02, Fig. 2C; ETC Occl 0.87 ± 0.02; Fig. 2E; for both, mixed model ANOVA nested on set, effect of treatment p<0.0001).

In summary, despite some mouse-to-mouse variability which is more marked for excitatory neurons, short-duration naris occlusion comparable to the sensory deprivation produced by a mild common cold (Fokkens et al., 2012), is effective overall in reducing activity levels in multiple OB cell types.

### Lack of structural and intrinsic activity-dependent plasticity in excitatory neurons

Previous *in vitro* work from our laboratory has demonstrated that both GABAergic and GABA-negative neurons in bulbar dissociated cultures respond to 24 h manipulations of neuronal activity by modulating the length and/or position of their AIS (Chand et al., 2015). This finding raised a number of questions, namely, (a) whether AIS plasticity also occurs *in vivo* in response to a sensory manipulation of similar duration, (b) if so, in which cell types, and, finally (c) whether structural plasticity at the AIS is accompanied by functional plasticity of the neurons’ intrinsic excitability.

In multiple cell types after 24 h naris occlusion, we performed *ex vivo* immunohistochemistry to quantify AIS position and length, and whole-cell patch clamp recording in acute slices to assess neurons’ passive and active electrophysiological properties.

In fixed slices of juvenile C57Bl6 mice, we identified M/TCs by staining the neurofilament protein H, clone SMI-32 (Ashwell, 2006). AISes were identified with staining against ankyrin-G (AnkG, Fig. 4A), and measured in 3D (see Methods). M/TCs all have a have prominent and reliably-oriented axon which arises directly from the soma and projects towards the granule cell layer of the OB. Their AnkG-positive AISs tend to be ~25 μm in length and proximally located (Lorincz and Nusser, 2008).

**Figure 4.**
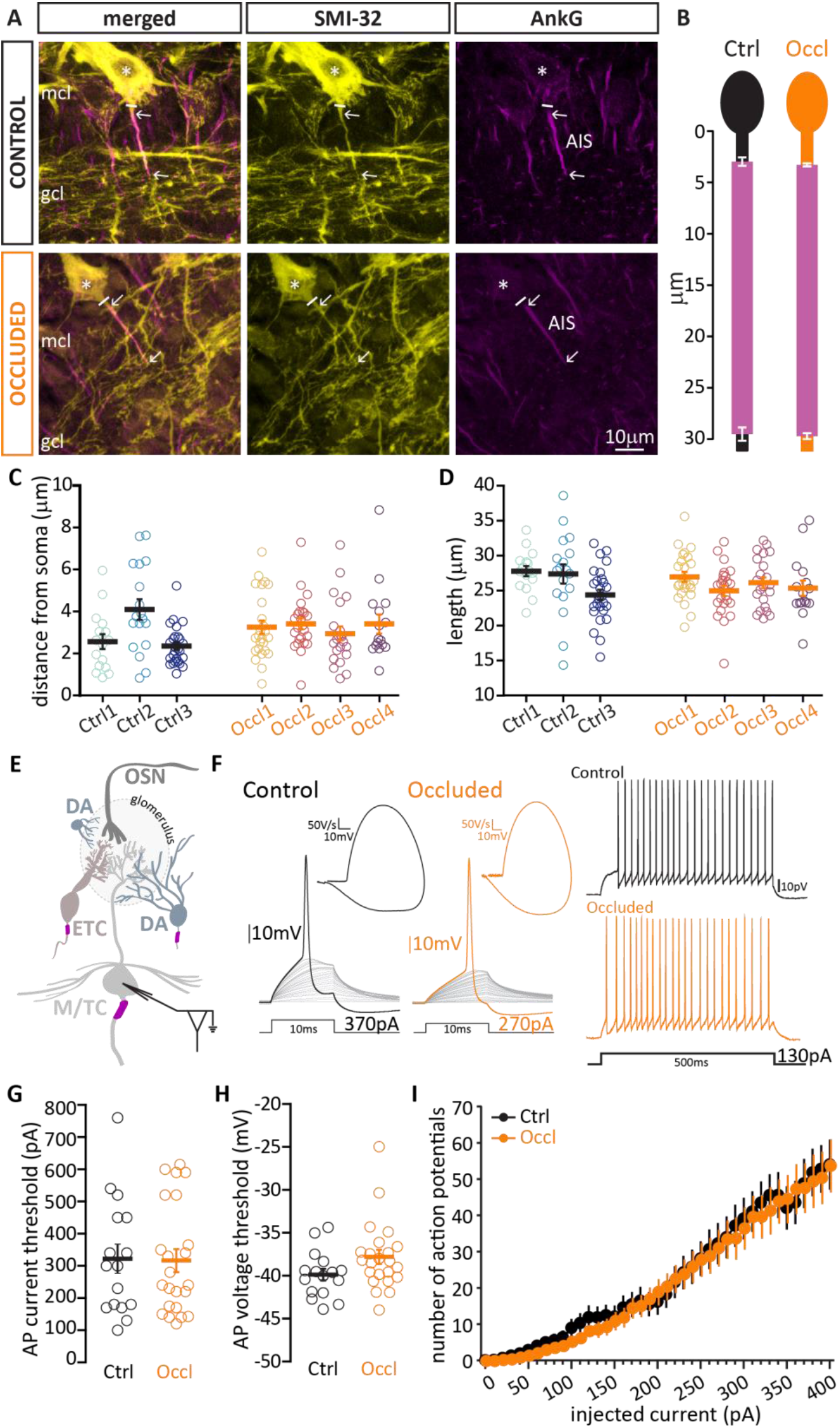
Brief unilateral naris occlusion fails to induce structural plasticity at the axon initial segment or plasticity of intrinsic excitability in mitral/tufted cells. **(A)** Example average intensity projection image of bulbar mitral/tufted cells (M/TCs, visualized via SMI-32 staining, yellow) and AIS marker ankyrin G (AnkG, magenta) in control and occluded mice. mcl = mitral cell layer; gcl = granule cells layer. The solid line indicates the emergence of the axonal process from the soma (asterisk); arrows indicate AIS start and end positions. **(B)** AIS plot shows the mean ± SEM start and end position for each group. **(C)** AIS distance from soma in M/TCs from control (N=3, n=61; mean±SEM 2.92 ± 0.21 μm) and occluded (N=4, n=87, 3.25 ± 0.15 μm) mice. Empty circles represent individual cells and different colours indicate different mice; thick line shows mean ± SEM. **(D)** Length of the same M/TCs AISes presented in C from control (mean±SEM 26.17 ± 0.58 μm) and occluded (25.91 ± 0.40 μm) mice. **(E)** Schematic representation of the experimental strategy for whole-cell recordings: acute 300μm OB slices were obtained from P21-35 mice, and M/TCs were targeted for whole-cell patchclamp recording based on soma location. **(F)** Left: example current-clamp traces of single APs fired to threshold 10ms somatic current injection by control (black) and occluded (orange) M/TCs, and their associated phase plane plots. Right: Example current-clamp traces of multiple APs fired in response to a 130pA/500 ms somatic current injection in control and occluded cells. **(G)** Single action potential current threshold in control (mean±SEM 322.5 ± 45.37 pA) and occluded (316.90 ± 35.71 pA) M/T cells. **(H)** Single action potential voltage threshold in control (mean±SEM −39.86 ± 0.67 mV) and occluded (−37.80 ± 0.84 mV) M/TCs. **(I)** Input-output curve of 500ms-duration current injection magnitude versus mean ± SEM spike number for each group. Empty circles represent individual cells (Ctrl, n=16; Occl; n=23); thick line and full circles show mean ± SEM.

We found no difference in AIS distance from the soma (Ctrl, N=3, n=61, mean ± SEM 2.92 ± 0.21 μm; Occl, N=4, n=87, 3.25 ± 0.15 mm; mixed model ANOVA nested on mouse, p=0.31), nor in AIS length (Ctrl, mean ± SEM 26.17 ± 0.58 mm; Occl, 25.91 ± 0.40 mm; mixed model ANOVA nested on mouse, p=0.64) between control and occluded M/TCs (Fig. 4B-D). This lack of structural AIS plasticity was mirrored by an equal absence of plastic changes in M/TCs’ intrinsic excitability. When probed with short current injections (10 ms, Fig. 4F right), control and occluded M/TCs fired an action potential at similar thresholds, both in terms of injected current (Fig. 4G; Ctrl, mean ± SEM 323 ± 45 pA; Occl, 317 ± 36 pA; unpaired t-test, p=0.92) and somatic membrane voltage (Fig. 4h; Ctrl, mean ± SEM – 39.86 ± 0.67 mV; Occl, −37.80 ± 0.83 mV; Mann-Whitney test, p=0.07). When probed with longer 500 ms current injections to elicit repetitive action potential firing (Fig. 4F left) we again found no difference between the two groups (Fig. 4I). Moreover, control and occluded M/TCs did not differ significantly in any other measured electrophysiological property, passive or active (Table 1).

**Table 1.** Intrinsic electrophysiological properties of mitral/tufted cells (M/TCs). Mean values ± SEM of passive, action potential and repetitive firing properties for control and occluded M/T cells. Statistical differences between groups were calculated with an unpaired t-test for normally-distributed data (“t”) or with a Mann–Whitney test for non-normally distributed data (“MW”). Individual data points and example traces are presented in Figure 4.

| Mitral/tufted cells |  |  |  |
| --- | --- | --- | --- |
| | Control<br>(mean ± SEM, [n]) | Occluded<br>(mean ± SEM, [n]) | Test type,<br>*p*-value |
| **PASSIVE PROPERTIES** |  |  |  |
| Membrane capacitance (pF) | 66 ± 4, [24] | 65 ± 4, [26] | t, 0.77 |
| Resting membrane potential (mV) | -49.12 ± 1.318, [4] | -51.33 ± 1.535, [8] | t, 0.41 |
| Input Resistance (mΩ) | 135 ± 19, [24] | 108 ± 11, [26] | MW, 0.33 |
| **ACTION POTENTIAL PROPERTIES** |  |  |  |
| Threshold (pA/pF) | 323 ± 45, [16] | 317 ± 36, [23] | t, 0.92 |
| Threshold (mV) | -39.86 ± 0.67, [16] | -37.80 ± 0.83, [23] | MW, 0.07 |
| Max voltage reached (mV) | 29.29 ± 1.64, [16] | 30.81 ± 1.21, [23] | t, 0.45 |
| Peak amplitude (mV) | 69.15 ± 1.38, [16] | 68.61 ± 1.28, [23] | t, 0.78 |
| Width at half-height (ms) | 0.41 ± 0.02, [15] | 0.45 ± 0.02, [23] | t, 0.15 |
| Rate of rise (max $dV/dt$, mV*ms) | 366 ± 16, [16] | 346 ± 12, [23] | t, 0.32 |
| Onset rapidness (1/ms) | 33.08 ± 1.16, [16] | 32.55 ± 0.95, [23] | t, 0.72 |
| After hyper polarization (AHP, mV) | -54.12 ± 0.85, [22] | -53.88 ± 0.59, [24] | t, 0.82 |
| AHP relative to threshold (mV) | 16.24 ± 0.74, [22] | 18.16 ± 0.80, [24] | t, 0.09 |
| **REPETITIVE FIRING PROPERTIES** |  |  |  |
| Rheobase (pA) | 108 ± 17.72, [22] | 122.2 ± 16.06, [25] | t, 0.55 |
| Max number of action potentials | 64.39 ± 6.33, [22] | 58.75 ± 6.16, [25] | t, 0.53 |
| First action potential delay (ms) | 392 ± 16, [22] | 368 ± 18, [25] | t, 0.33 |
| Inter-spike interval CV | 0.48 ± 0.07, [22] | 0.38 ± 0.05, [26] | MW, 0.44 |

Similarly, we also found no evidence for structural or intrinsic activity-dependent plasticity in ETCs. In these experiments we visualized ETCs in fixed tissue by staining for cholecystokinin (CCK, Fig. 4A; (Liu and Shipley, 1994)). We found that, as for M/TCs, ETC AISes are prominent AnkG-positive segments located quite proximally on a process originating directly from the soma. These AISes were equally distant from the soma (Fig. 5C; Ctrl, N=3, n=65, mean ± SEM 2.67 ± 0.23 μm; Occl, N=3, n=62, 2.496 ± 0.22 μm; mixed model ANOVA nested on mouse, p=0.90) and equally long (Fig. 5D; Ctrl, mean ± SEM 18.52 ± 0.39 μm; Occl, 19.94 ± 0.59 μm; mixed model ANOVA nested on mouse, p=0.18) in control and occluded mice. Moreover, when probed electrophysiologically in acute slices (Fig. 5F-G), ETCs from control and occluded mice fired single action potentials at similar thresholds (current threshold, Fig. 5G, Ctrl, mean ± SEM 103 ± 8 pA; Occl, 112 ± 7 pA, Mann-Whitney test, p=0.50; voltage threshold, Fig. 5H, Ctrl −39.10 ± 0.48 mV; Occl −38.56 ± 0.48 mV; Mann-Whitney test, p=0.57), and similarly modulated their repetitive firing in response to long current injections of increasing intensity (Fig. 5I, Table 2).

**Figure 5.**
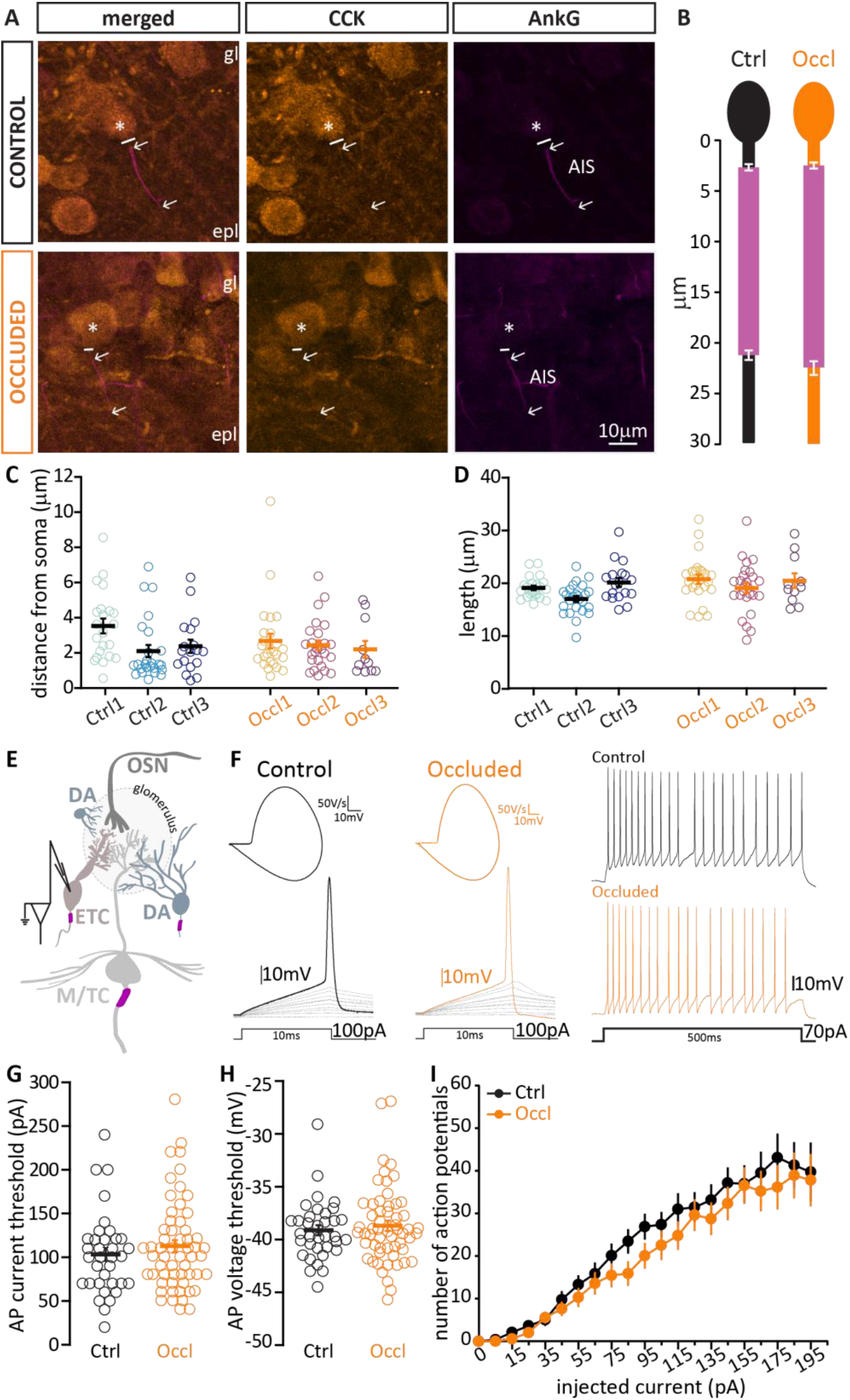
Brief unilateral naris occlusion fails to induce structural plasticity at the axon initial segment or plasticity of intrinsic excitability in external tufted cells. (A) Example average intensity projection image of bulbar external tufted cells (ETCs, visualized via staining against cholecystokinin, CCK, amber) and AIS marker ankyrin-G (AnkG, magenta) in control and occluded mice. gl = glomerular layer; epl = external plexiform layer. The solid line indicates the emergence of the axonal process from the soma (asterisk); arrows indicate AIS start and end positions. **(B)** AIS plot shows the mean ± SEM start and end position for each group. **(C)** AIS distance from soma in ETCs from control (N=3, n=65; mean+SEM 2.67 ± 0.23 μm) and occluded (N=3, n=62, 2.496 ± 0.22 μm) mice. Empty circles represent individual cells and different colours indicate different mice; thick line shows mean ± SEM. **(D)** Length of the same ETC AISes presented in (C) from control (mean±SEM 18.52 ± 0.39 μm) and occluded (19.94 ± 0.59 μm) mice. **(E)** Schematic representation of the experimental strategy for whole-cell recordings: acute 300 μm OB slices were obtained from P21-35 mice, and ETCs were targeted for whole-cell patch-clamp recording based on location and soma size (see methods for inclusion criteria). **(F)** Left: example current-clamp traces of single APs fired to threshold 10ms somatic current injection by control (black) and occluded (orange) ETCs, and their phase plane plots. Right: Example current-clamp traces of multiple APs fired in response to a 70pA/500 ms somatic current injection in control and occluded cells. **(G)** Single action potential current threshold in control (mean±SEM 103 ± 8 pA) and occluded (112 ± 7 pA) ETCs. **(H)** Single action potential voltage threshold in control (mean±SEM −39.10 ± 0.48 mV) and occluded (−38.56 ± 0.48 mV) ETCs. **(I)** Input-output curve of injected 500ms-long currents versus mean ± SEM spike number for each group. Empty circles represent individual cells (Ctrl, n=35; Occl; n=57); thick line and full circles shows mean ± SEM.

**Table 2.** Intrinsic electrophysiological properties of external tufted cells (ETCs) Mean values ± SEM of passive, action potential and repetitive firing properties for control and occluded ET cells. Statistical differences between groups were calculated with an unpaired t-test for normally-distributed data (“t”) or with a Mann–Whitney test for non-normally distributed data (“MW”). Individual data points and example traces are presented in Figure 5.

| External tufted cells |  |  |  |
| --- | --- | --- | --- |
| | Control<br>(mean ± SEM, [n]) | Occluded<br>(mean ± SEM, [n]) | Test type,<br>*p*-value |
| **PASSIVE PROPERTIES** |  |  |  |
| Membrane capacitance (pF) | 43.43 ± 2.13, [35] | 41.92 ± 1.67, [57] | t, 0.58 |
| Resting membrane potential (mV) | -57.33 ± 0.92, [36] | -56.58 ± 0.94, [64] | t, 0.60 |
| Input Resistance (mΩ) | 271 ± 27, [35] | 242 ± 17, [57] | MW, 0.29 |
| **ACTION POTENTIAL PROPERTIES** |  |  |  |
| Threshold (pA/pF) | 103 ± 8, [35] | 112 ± 7, [57] | MW, 0.50 |
| Threshold (mV) | -39.10 ± 0.48, [35] | -38.56 ± 0.48, [57] | MW, 0.57 |
| Max voltage reached (mV) | 22.08 ± 1.18, [35] | 22.69 ± 0.69, [57] | t, 0.64 |
| Peak amplitude (mV) | 61.17 ± 1.14, [35] | 61.26 ± 0.82, [57] | t, 0.94 |
| Width at half-height (ms) | 0.53 ± 0.02, [35] | 0.52 ± 0.01, [57] | MW, 0.94 |
| Rate of rise (max $dV/dt$, mV*ms) | 205 ± 7, [35] | 211 ± 4, [57] | t, 0.44 |
| Onset rapidness (1/ms) | 29.65 ± 0.85, [30] | 30.18 ± 0.94, [37] | MW, 0.42 |
| After hyper polarization (AHP, mV) | -52.04 ± 0.45, [23] | -52.30 ± 0.60, [38] | t, 0.76 |
| AHP relative to threshold (mV) | 15.41 ± 0.85, [23] | 17.14 ± 0.58, [38] | t, 0.09 |
| **REPETITIVE FIRING PROPERTIES** |  |  |  |
| Rheobase (pA) | 41 ± 8 pA, [23] | 45 ± 7 pA, [38] | MW, 0.73 |
| Max number of action potentials | 61 ± 4, [30] | 56 ± 3, [45] | t, 0.34 |
| First action potential delay (ms) | 187 ± 23, [23] | 171 ± 22, [38] | t, 0.63 |
| Inter-spike interval CV | 0.33 ± 0.04, [30] | 0.38 ± 0.04, [45] | MW, 0.65 |

Taken together, these results confirm that while both major classes of bulbar excitatory neurons experience an overall drop in activity after 24 h sensory deprivation (Fig. 2), they do not respond by altering the structural features of their AIS or their intrinsic physiological properties.

### Both inhibitory dopaminergic neuron subclasses downregulate their TH expression levels in response to brief naris occlusion

In other brain areas inhibitory interneurons can act as first responders in the early phases of adaptation to changed incoming activity, plastically changing their overall structure and function to maintain circuit homeostasis (Gainey and Feldman, 2017; Hartmann et al., 2008; Keck et al., 2017; Knott et al., 2002; Yin and Yuan, 2014). Given the lack of plasticity in glutamatergic OB neurons following brief 24 h naris occlusion, we reasoned that plastic responses might therefore be more evident in OB inhibitory interneurons. Because of their renowned plasticity *in vivo* and their ability to undergo activity-dependent AIS changes *in vitro* (Bonzano et al., 2016; Chand et al., 2015), we focused on the bulb’s DA population to address this question.

Bulbar DA neurons are unique amongst other glomerular layer inhibitory neurons because of their well-described plasticity in neurotransmitter-synthesising enzyme expression. Changes in sensory input, including those induced by unilateral naris occlusion, are known to produce alterations in tyrosine hydroxylase (TH) expression at both the protein and mRNA levels (Baker et al., 1993; Cummings and Brunjes, 1997; Kosaka et al., 1987; Nadi et al., 1981). As with other forms of experience-dependent plasticity, these changes have been mostly investigated using long-duration manipulations. However, 2 days of deprivation were reported to induce a small, but significant, decrease in whole-bulb *Th* mRNA (Cho et al., 1996), whilst just one day of elevated activity was sufficient to increase TH immunofluorescence intensity or TH-GFP transgene expression, respectively, in dissociated and slice culture preparations (Akiba et al., 2007; Chand et al., 2015). We therefore set out to assess whether 24 h naris occlusion is sufficient to produce activity-dependent changes in TH expression *in vivo*, and if so whether these changes are observed in both axon-bearing and anaxonic OB DA subtypes.

In three sets of co-embedded control and occluded coronal bulbar slices (Fig. 2B) stained with an antibody against TH (Fig. 6A), we first confirmed that the overall density of labelled DA cells was unaffected by brief sensory deprivation (Fig. 6B; Ctrl mean ± SEM 42317 ± 3661 cells/mm^3^, Occl 39993 ± 4243 cells/mm^3^; unpaired t-test, p=0.68). In the knowledge that we were labelling a similar number of TH-positive cells in both groups, we then analyzed relative TH immunofluorescence levels in each set, normalizing the intensity of staining to average control values (see Methods). Given the inter-set variability noted in our cFos data (Fig 2), it was unsurprising to also observe such variability in relative TH intensity levels, especially for the smaller sample of much rarer large neurons (Fig. 6c-e). Nevertheless, multilevel analyses accounting for this variation revealed highly significant overall reductions in TH immunofluorescence levels in all DA cell groups – significant changes were observed in all DA cells (Fig. 6C), small putative anaxonic cells (Fig. 6D), and large putative axonal cells (Fig. 6E; mixed model ANOVA nested on set, all p<0.001). We further confirmed this latter phenotype in a smaller subset of DA cells with definitively identified AISes (Fig. 6F, AnkG in magenta; normalized TH intensity: Ctrl mean ± SEM 1.79 ± 0.13, Occl 1.37 ± 0.14, unpaired t–test, p=0.0498). We also found significant positive correlations between normalized TH and normalized cFos (Fig. 2) intensities for all groups. These were stronger for control neurons (norm TH vs. norm cFos, Ctrl small cells r^2^=0.48, big cells r^2^=0.80; Occl small cells r^2^=0.27, big cells r^2^=0.18; all p<0.02), suggesting that the mechanisms leading to activity-dependent TH and cFos changes in individual OB DA neurons are only loosely coupled. Overall, given that alterations in OB TH levels are often used to confirm the effectiveness of olfactory sensory manipulations (Cockerham et al., 2009; Grier et al., 2016; Kass et al., 2013), these data supplement the immediate early gene analysis (Fig. 2) to show that 24 h naris occlusion strongly and reliably downregulates activity in both subclasses of OB DA interneurons. They also provide evidence for, to date, the fastest activity-dependent TH change observed in this cell class *in vivo* (Byrne, 2019)

**Figure 6.**
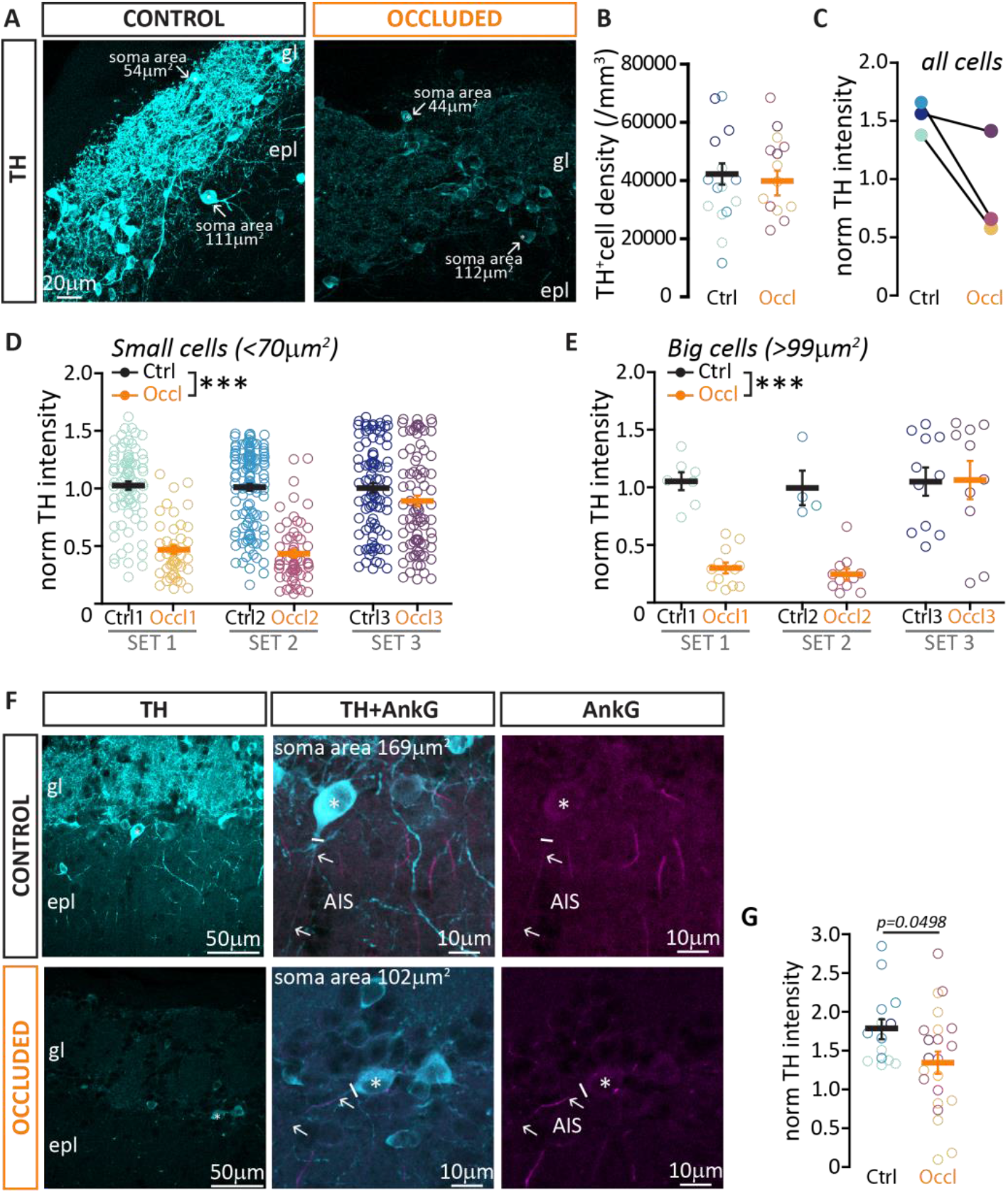
Brief unilateral naris occlusion decreases the expression of tyrosine hydroxylase in both DA subtypes. **(A)** Example maximum intensity projection image of dopaminergic neurons (DA cells, cyan) visualized via anti-tyrosine hydroxylase (TH) immunolabel in control (black) and occluded (orange) mice. gl = glomerular layer; epl = external plexiform layer. Arrows indicate TH positive DA cells representative of the two subtypes when defined by soma area. **(B)** Average density of TH-positive cells (of any soma size) in control (mean±SEM 42317 ± 3661) and occluded (39993 ± 4243) glomerular layer. Empty circles represent individual image stacks (Ctrl, n=14; Occl; n=15) and different colours indicate different mice; thick lines show mean ± SEM. **(C)** TH intensity in DA cells of any soma size, normalized to co-embedded control measures, in three sets of control + occluded OB slices. **(D)** Normalized TH intensity of DA cells with soma size < 70 μm^2^ (putative anaxonic cells), from 3 sets of Ctrl (black, n=298, overall mean±SEM 1.01 ± 0.02) and Occl (orange, n=192, 0.64 ± 0.03) mice. **(E)** Normalized TH intensity of DA cells with soma size > 99 μm^2^ (putative axon-bearing DA cells), from 3 sets of Ctrl (black, n=22, overall mean±SEM 1.01 ± 0.07) and Occl (orange, n=33, 0.51 ± 0.08) mice. **(F)** Example average intensity projection images of TH expression (cyan) in DA cells with an identified ankyrin-G (AnkG; magenta)-positive AIS (arrows). The solid line indicates the emergence of the axonal process from the soma (asterisk). gl = glomerular layer; epl = external plexiform layer. **(G)** Normalized TH intensity in AnkG-positive DA cells in control (n=14, mean±SEM 1.79 ± 0.13) and occluded (n=22,1.37 ± 0.14) mice. Empty circles represent individual cells and different colours indicate different mice (n=3, N=3 for both Ctrl and Occl); thick lines shows mean ± SEM; *= p<0.05, ***= p<0.001.

### Anaxonic DA neurons do not modulate their intrinsic excitably following brief sensory deprivation

The vast majority of DA neurons are anaxonic cells (Galliano et al., 2018), which by locally releasing GABA and dopamine in the glomerular layer help to control the overall gain of OSN➔M/TC transmission (McGann, 2013; Vaaga et al., 2017). Highly plastic, they retain the capability to regenerate throughout life (Bonzano et al., 2016; De Marchis et al., 2007; Galliano et al., 2018; Lledo et al., 2006). However, although they regulate their levels of TH expression in response to 24 h naris occlusion (Fig 6), we found that the same manipulation did not change their intrinsic excitability.

We performed whole-cell patch clamp recordings in control and occluded DAT-tdTomato mice (Bäckman et al., 2006; Madisen et al., 2010). This transgenic labelling approach produces red fluorescent tdT-positive glomerular layer cells that are ^~^75-85% co-labelled for TH (Fig. 7A; (Byrne, 2019; Galliano et al., 2018; Vaaga et al., 2017)). The remaining tdT-positive/TH-negative non-dopaminergic labelled OB neurons in these mice are of the calretinin-expressing OB interneuron class and can be readily identified by their unique physiological properties (Byrne, 2019; Pignatelli et al., 2005; Sanz Diez et al., 2019), so these were excluded from our analyses. Anaxonic DA cells, which are over-represented in DAT-tdT mice (Galliano et al., 2018), were functionally classified by assessing the nature of their action potential phase plane plot of single spikes fired in response to 10 ms somatic current injection (Fig. 7B). A smooth, monophasic phase plane plot is indicative of AP initiation at the somatic recording site, and can be used as a proxy indicator of anaxonic morphology. In contrast, a distinctive biphasic phase plane plot waveform indicates that the AP initiated at a non-somatic location – usually the AIS – and can be used as a proxy for axon-bearing identity (Bean, 2007; Bender and Trussell, 2012; Chand et al., 2015; Coombs et al., 1957; Foust et al., 2010; Galliano et al., 2018; Jenerick, 1963; Khaliq et al., 2003; Kole et al., 2007; Shu et al., 2007). Indeed, we confirmed that monophasic, putative anaxonic cells had smaller soma sizes than putative axon-bearing neurons with biphasic phase plane plot signatures (see below; monophasic cells mean ± SEM 56.36 ± 3.40 μm^2^; biphasic cells 89.44 ± 5.19 μm^2^; unpaired t-test, p<0.001)(Chand et al., 2015; Galliano et al., 2018).

**Figure 7.**
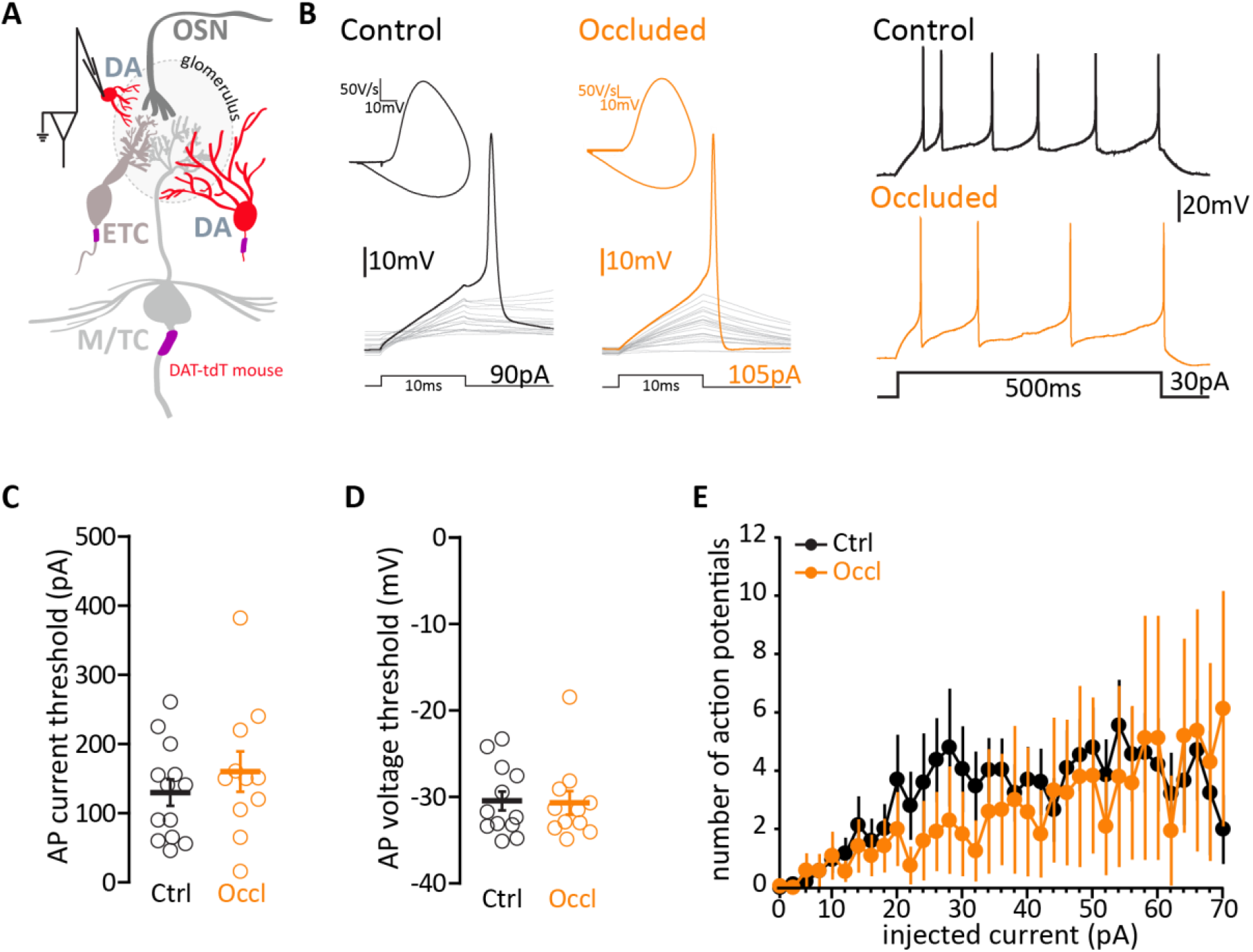
Brief unilateral naris occlusion does not alter the intrinsic excitability of monophasic/putative anaxonic DA cells. **(A)** Schematic representation of the experimental strategy for whole-cell recordings: acute 300 μm OB slices were obtained from P21-35 DAT-tdTomato mice, and monophasic DA cells were targeted for whole-cell patch-clamp recordings based on red fluorescence and soma size. **(B)** Left: example current-clamp traces of single APs fired to threshold 10 ms somatic current injection by control (black) and occluded (orange) DA cells, and their associated monophasic phase plane plots. Right: Example current-clamp traces of multiple APs fired in response to a 30 pA / 500 ms somatic current injection in control and occluded cells. **(C)** Single action potential current threshold in control (mean±SEM 130 ± 20 pA) and occluded (160 ± 30 pA) monophasic DA neurons. **(D)** Single action potential voltage threshold in control (mean±SEM −30.47 ± 1.09 mV) and occluded (−30.70 ± 1.37 mV) monophasic DA neurons. **(E)** Input-output curve of 500 ms-duration current injection magnitude versus mean ± SEM spike number for each group. Empty circles represent individual cells (Ctrl, n=13; Occl; n=11); thick line and full circles show mean ± SEM.

We found that, while sitting at a more depolarized resting membrane potential than their control counterparts, monophasic/putative-anaxonic DA cells from occluded mice showed no other significant differences in their passive membrane properties (Table 3). Measures of intrinsic excitability – importantly measured from identical baseline voltage – were indistinguishable between the two groups. Control and occluded monophasic neurons fired single spikes at similar thresholds (current threshold, Fig. 7D, Ctrl, mean ± SEM 129.7 ± 19.2 pA; Occl, 160 ± 29.23 pA, unpaired t-test, p=0.38; voltage threshold, Fig. 7D, Ctrl −30.47 ± 1.09 mV; Occl −30.70 ± 1.37 mV; Mann-Whitney test, p=0.86), and, when probed with longer current injections of increasing intensity, fired similar numbers of multiple action potentials (Fig. 7E, Table 3).

**Table 3.** Intrinsic electrophysiological properties of monophasic/putative anaxonic DA cells. Mean values ± SEM of passive, action potential and repetitive firing properties for control and occluded monophasic/putative anaxonic DA cells. Statistical differences between groups were calculated with an unpaired t-test for normally-distributed data (“t”; with Welch’s correction “t(W)”) or with a Mann–Whitney test for non-normally distributed data (“MW”). Grey shading indicates statistically significant difference. Individual data points and example traces are presented in Figure 7.

| Monophasic dopaminergic cells (putative anaxonic) |  |  |  |
| --- | --- | --- | --- |
| | Control<br>(mean ± SEM, [n]) | Occluded<br>(mean ± SEM, [n]) | Test type,<br>*p*-value |
| **PASSIVE PROPERTIES** |  |  |  |
| Membrane capacitance (pF) | 19.17 ± 2.18, [15] | 20.81 ± 1.77, [12] | MW, 0.37 |
| Resting membrane potential (mV) | -77.87 ± 1.92, [15] | -70.50 ± 2.49, [12] | t, 0.03 |
| Input Resistance (mΩ) | 960 ± 272, [15] | 694 ± 223, [12] | MW, 0.21 |
| **ACTION POTENTIAL PROPERTIES** |  |  |  |
| Threshold (pA) | 129.7 ± 19.2, [13] | 160 ± 29.23, [11] | t, 0.38 |
| Threshold (mV) | -30.47 ± 1.09, [13] | -30.70 ± 1.37, [11] | MW, 0.86 |
| Max voltage reached (mV) | 19.55 ± 2.42, [13] | 23.19 ± 1.78, [11] | t, 0.26 |
| Peak amplitude (mV) | 50.01 ± 2.35, [13] | 53.89 ± 2.89, [11] | MW, 0.22 |
| Width at half-height (ms) | 0.54 ± 0.03, [13] | 0.55 ± 0.03, [11] | t, 0.80 |
| Rate of rise (max $dV/dt$, mV*ms) | 240.7 ± 15.82, [13] | 254.8 ± 19.63, [11] | t, 0.58 |
| Onset rapidness (1/ms) | 3.94 ± 0.29, [13] | 3.23 ± 0.20, [11] | t, 0.06 |
| After hyper polarization (AHP, mV) | -54.39 ± 1.44, [14] | -54.83 ± 1.34, [12] | t, 0.83 |
| AHP relative to threshold (mV) | 24.58 ± 1.27, [14] | 25.87 ± 1.49, [12] | t, 0.51 |
| **REPETITIVE FIRING PROPERTIES** |  |  |  |
| Rheobase (pA) | 61 ± 19, [14] | 86 ± 20, [12] | t, 0.23 |
| Max number of action potentials | 10 ± 2, [14] | 7 ± 2, [12] | MW, 0.16 |
| First action potential delay (ms) | 169.2 ± 38.99, [14] | 91.34 ± 21.92, [12] | t(W), 0.10 |
| Inter-spike interval CV | 0.28 ± 0.04, [14] | 0.26 ± 0.04, [11] | t, 0.72 |

Overall, in putative anaxonic/monophasic DA cells the decreases in c-fos and TH expression observed after 24 h naris occlusion are not accompanied by any significant alterations in intrinsic excitability.

### DA cells equipped with an axon shorten their axon initial segment and decrease their intrinsic excitability in response to 24 h naris occlusion

Far less abundant than their anaxonic neighbours, axon-bearing DA neurons tend to have a large soma, and dendrites that branch more widely within the glomerular layer (Galliano et al., 2018). Similarly to anaxonic DA cells, they respond to 24 h naris occlusion by decreasing cFos and TH expression (Figs. 3 and 6), but they lack a key characteristic of the former: the dramatic whole-cell structural plasticity which is the ability to regenerate throughout life. Instead of undergoing lifelong neurogenesis, axon-bearing OB DA cells are exclusively born during early embryonic stages (Galliano et al., 2018). However, we have previously shown that, *in vitro*, this DA subtype can undergo a much subtler type of structural plasticity in the form of AIS alterations. In particular, 24 h reduced activity in the presence of tetrodotoxin was associated with decreased AIS length in this cell type (Chand et al., 2015). We therefore set out to investigate whether similar AIS plasticity also occurs *in vivo* in response to the same duration of sensory deprivation.

As for AIS analysis in excitatory neurons, we performed immunohistochemistry in fixed slices of juvenile C57Bl6 mice, double stained for TH to identify DA neurons (Fig. 8A, cyan) and ankyrin-G to measure AISes (AnkG, Fig. 8A, magenta). A current leitmotiv in the biology of DA neurons is their striking heterogeneity (Chand et al., 2015; Henny et al., 2012; Kosaka et al., 2019; Morales and Margolis, 2017; Romanov et al., 2017; Zhang et al., 2007), and in OB DA cells here this was also evident in the structure and location of their AIS. We found that OB AISes are of reasonably consistent length (coefficient of variation, CV = 0.34 in control cells), but can be situated at highly variable distances from the soma (control CV = 0.75). Contrary to findings in midbrain DA cells (González-Cabrera et al., 2017; Meza et al., 2018) and in OB dissociated cultures (Chand et al., 2015) we found no consistent relationship between these parameters in bulbar DA neurons (Spearmen coefficient of AIS length vs soma distance: Ctrl r=0.03, Occl, r=0.04; both p>0.73). We also noted that that the AIS of an OB DA neuron can be located either on a process that directly emanates from the soma (“soma-origin” AIS), or on a process separated from the soma by one or more branch nodes (“dendrite-origin” AIS, Ctrl n=31, Occl n=33; Fig. 8A-B)(González-Cabrera et al., 2017; Höfflin et al., 2017; Houston et al., 2017; Kosaka et al., 2019; Thome et al., 2014; Yang et al., 2019). While this peculiar axonal arrangement challenges the traditional view on neuronal input-output transformation (Kaifosh and Losonczy, 2014), it is not unique to bulbar DA neurons. Indeed, midbrain DA neurons have been shown to carry “dendrite-origin” AISes (González-Cabrera et al., 2017; Yang et al., 2019), and recently the overall variability in AIS length and location in these neurons has been proposed to play a key role in the maintenance of an appropriate pacemaking rhythm in the context of variable dendritic branching (Moubarak et al., 2019). Moreover, “dendrite-origin” AISes are not exclusive to DA neurons: common in invertebrates (Triarhou, 2014), they have also been described in cat and mouse cortex (Hamada et al., 2016; Höfflin et al., 2017; Meyer and Wahle, 1988), in hippocampal pyramidal cells (Thome et al., 2014), and in cerebellar granule cells (Houston et al., 2017).

**Figure 8.**
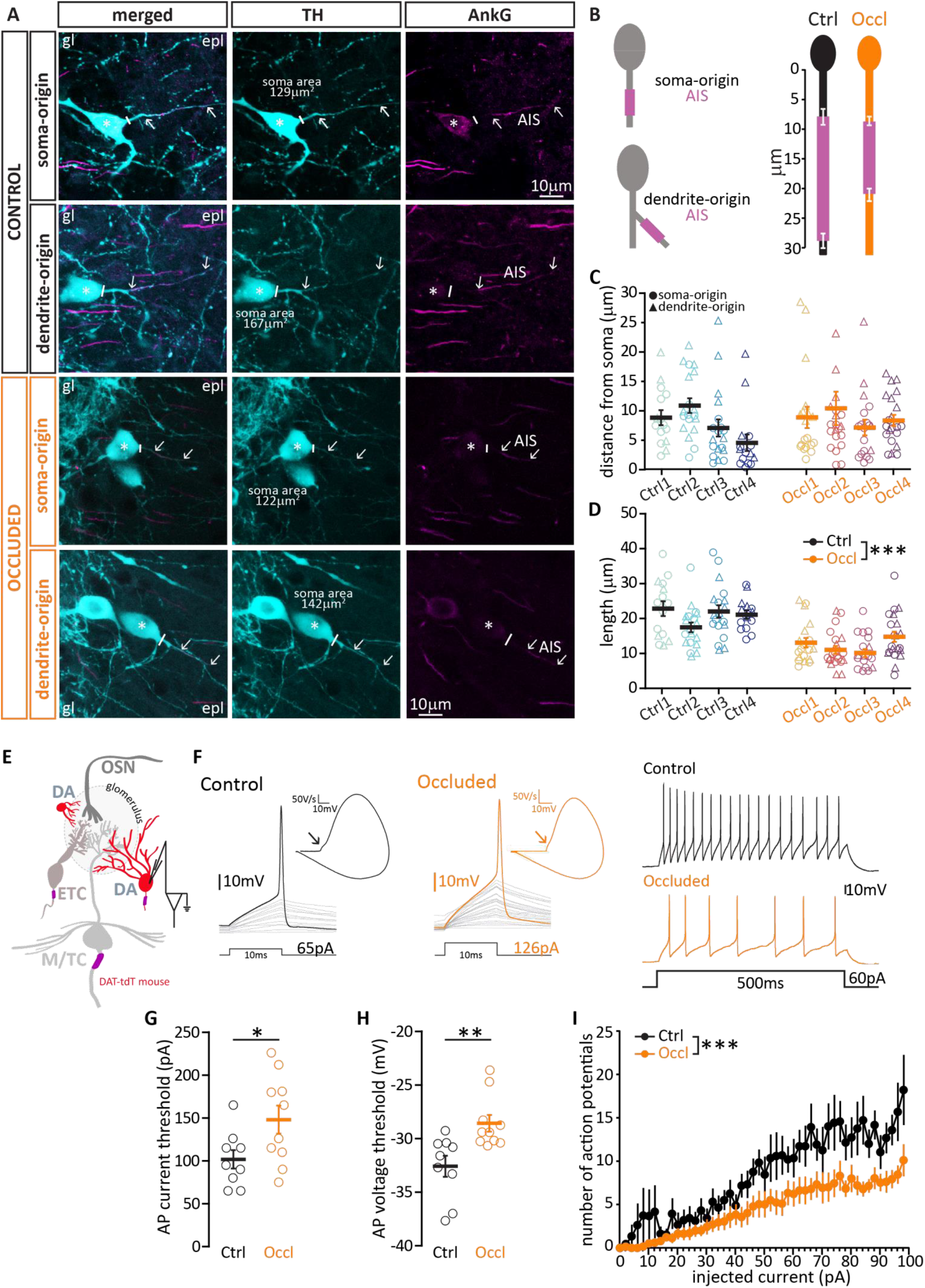
Brief unilateral naris occlusion results in shorter axon initial segments and decreased intrinsic excitability in biphasic/putative axon-bearing DA cells. **(A)** Example average intensity projections images of bulbar axonbearing DA cells, visualized via staining for tyrosine hydroxylase (TH, cyan) and the AIS marker ankyrin-G (AnkG, magenta) in control and occluded mice. In DA neurons, AISes can be found either on a process originating directly from the soma (soma-origin), or on a process separated from the soma by one or more nodes (dendrite-origin). gl = glomerular layer; epl = exernal plexiform layer. The solid line indicates the emergence of the axonal process from the soma (asterisk); arrows indicate AIS start and end positions. **(B)** Left: schematic representation of soma-origin and dendrite-origin AISs. Right: Average AIS plots (soma+dendrite origin) show the mean ± SEM AIS start and end positions for each group. **(C)** AIS distance from soma in DA cells from control (N=4, n=68; mean±SEM 7.91 ± 0.73 μm) and occluded (N=4, n=80, 8.47 ± 0.94 μm) mice. For representation clarity, one outlier for distance from soma (62 μm, occluded group) is not included in the figure, but is included in all averages and analysis. Empty symbols represent individual cells and different colours indicate different mice; circles indicate soma-origin AIS, triangles indicate dendrite-origin AIS; thick line shows mean ± SEM. **(D)** Length of the same DA AISes presented in (C) from control (mean±SEM 20.74 ± 0.84 μm) and occluded (12.29 ± 0.66 μm) mice. **(E)** Schematic representation of the experimental strategy for whole-cell recordings: acute 300 μm OB slices were obtained from P21-35 DAT-tdTomato mice, and putative axon-bearing DA cells were targeted for whole-cell patch-clamp recordings based on red fluorescence and soma size. **(F)** Left: example current-clamp traces of single APs fired to threshold 10ms somatic current injection by control (black) and occluded (orange) DA cells, and their associated biphasic plane plots. Arrow points to the AIS-dependent first action potential phase. Right: Example current-clamp traces of multiple APs fired in response to a 60 pA / 500 ms somatic current injection in control and occluded cells. **(G)** Single action potential current threshold in control (mean±SEM 102 ± 11 pA) and occluded (148 ± 16 pA) biphasic DA cells. **(H)** Single action potential voltage threshold in control (mean±SEM −32.58 ± 0.99 mV) and occluded (−28.57 ± 0.78 mV) biphasic DA cells. **(I)** Inputoutput curve of 500 ms-duration injected current magnitude versus mean ± SEM spike number for each group. Empty circles represent individual cells (Ctrl, n=9; Occl; n=10); thick line and full circles show mean ± SEM; *= p<0.05, **= p<0.01, ***= p<0.001.

Occlusion did not affect the proportion of soma-vs. dendrite-origin AISes amongst the OB DA axon-bearing population (Soma: Ctrl n=37, Occl n=47; Dendrite: Ctrl n=31, Occl n=33; FET for proportions Ctrl vs Occl, p=0.62), nor did it affect the distance of the AIS start position from the soma, independent of axon origin (Fig. 8C, note different symbols to indicate axon origin; Ctrl, mean ± SEM 7.91 ± 0.73 μm; Occl, 8.47 ± 0.94 μm; mixed model ANOVA nested on mouse, effect of treatment p = 0.19; effect of axon origin, p = 0.42; effect of interaction, p = 0.17). We did, however, find a sizeable and consistent activity-dependent difference in AIS length, with AISes in occluded DA neurons being significantly shorter than those in control cells (Fig. 8D; Ctrl, mean ± SEM 20.74 ± 0.84 μm; Occluded 12.29 ± 0.66 μm; mixed model ANOVA nested on mouse, effect of treatment p < 0.0001; effect of axon origin, p = 0.39; effect of interaction, p = 0.49).

One key function of the AIS, which houses voltage-activated sodium channels at high density, is to initiate action potentials (Kole et al., 2007). Previous experimental evidence (*e.g.*, (Evans et al., 2015; Kuba et al., 2010)) and computational models (see, *e.g.*, (Goethals and Brette, 2020; Gulledge and Bravo, 2016; Hamada et al., 2016)) have shown that alterations in AIS length, all else being equal, are associated with decreases in neuronal excitability. So does the experience-dependent decrease in AIS length we observe in axon-bearing DA cells correlate with a reduced ability to fire action potentials? To test this prediction, we again turned to whole-cell patch clamp recordings in DAT-tdTomato mice, but this time we targeted red cells with a large soma (Fig. 8E), and used the biphasic nature of their action potential phase plots as a proxy for the presence of an AIS (Bean, 2007; Chand et al., 2015; Galliano et al., 2018). We found that, with no difference in key passive properties such as resting membrane potential and membrane resistance (Table 4), putative axon-bearing/biphasic DA cells recorded in acute slices obtained from occluded mice needed more current to reach threshold to generate an action potential (Fig. 8G, Ctrl, mean ± SEM 102 ± 11; Occl, 148 ± 16; unpaired t-test, p=0.04), and they did so at a more depolarized membrane voltage (Fig. 8H, Ctrl, mean ± SEM −32.58 ± 0.99; Occl, −28.57 ± 0.78; unpaired t-test, p=0.005). Moreover, when challenged with 500ms-long current injections of increasing amplitude, occluded DA cells fired less action potentials overall than control DA cells (Fig. 8I, mixed model ANOVA, effect of treatment, p=0.013; Table 4).

**Table 4.** Intrinsic electrophysiological properties of biphasic/putative axon-bearing DA cells. Mean values ± SEM of passive, action potential and repetitive firing properties for control and occluded biphasic/putative axon-bearing DA cells. Statistical differences between groups were calculated with an unpaired t-test for normally-distributed data (“t”) or with a Mann–Whitney test for non-normally distributed data (“MW”). Grey shading indicates statistically significant difference. Individual data points and example traces are presented in Figure 8.

| Biphasic dopaminergic cells (putative axon-bearing) |  |  |  |
| --- | --- | --- | --- |
| | Control<br>(mean ± SEM, [n]) | Occluded<br>(mean ± SEM, [n]) | Test type,<br>p-value |
| **PASSIVE PROPERTIES** |  |  |  |
| Membrane capacitance (pF) | 22.07 ± 2.21, [11] | 21.72 ± 2.07, [10] | t, 0.91 |
| Resting membrane potential (mV) | -74.27 ± 2.94, [11] | -77.50 ± 1.73, [10] | MW, 0.65 |
| Input Resistance (mΩ) | 573 ± 115, [11] | 631 ± 117, [10] | MW, 0.46 |
| **ACTION POTENTIAL PROPERTIES** |  |  |  |
| Threshold (pA) | 102 ± 11, [9] | 148 ± 16, [10] | t, 0.04 |
| Threshold (mV) | -32.58 ± 0.99, [9] | -28.57 ± 0.78, [10] | t, 0.005 |
| Max voltage reached (mV) | 17.61 ± 3.96, [9] | 19.87 ± 3.61, [10] | t, 0.68 |
| Peak amplitude (mV) | 50.17 ± 4.63, [9] | 48.43 ± 3.60, [10] | t, 0.76 |
| Width at half-height (ms) | 0.50 ± 0.04, [9] | 0.53 ± 0.03, [10] | t, 0.55 |
| Rate of rise (max $dV/dt$, mV*ms) | 250 ± 31, [9] | 227 ± 17, [10] | t, 0.51 |
| Onset rapidness (1/ms) | 8.22 ± 1.66, [9] | 6.63 ± 1.39, [10] | MW, 0.72 |
| After hyper polarization (AHP, mV) | -55.13 ± 1.50, [11] | -54.27 ± 2.71, [10] | MW, 0.55 |
| AHP relative to threshold (mV) | 24.46 ± 1.30, [11] | 25.17 ± 2.64, [10] | MW, 0.32 |
| **REPETITIVE FIRING PROPERTIES** |  |  |  |
| Rheobase (pA) | 32 ± 13, [11] | 25 ± 5, [10] | MW, 0.73 |
| Max number of action potentials | 21 ± 4, [11] | 15 ± 3, [10] | t, 0.23 |
| First action potential delay (ms) | 273 ± 45, [11] | 188 ± 50, [10] | t, 0.22 |
| Inter-spike interval CV | 0.24 ± 0.03, [10] | 0.22 ± 0.06, [9] | MW, 0.45 |

In summary, among the OB cell types we analyzed, axon-bearing DA interneurons are the only group that respond to brief, naturally-relevant sensory deprivation with a combination of biochemical (Fig. 6G), morphological (Fig. 8D) and functional (Fig. 8G-I) plastic changes.

## DISCUSSION

Our results demonstrate that, in young adult mice, brief 24 h sensory deprivation via the unilateral insertion of a custom-made naris plug is minimally-invasive yet sufficient to downregulate activity in olfactory bulb circuits. In response to this naturally-relevant manipulation (Fokkens et al., 2012) we find that only a very specific subtype of local inhibitory interneurons – axon-bearing DA cells located in the glomerular layer – respond with activity-dependent structural plasticity at their AIS and co-incident changes in their intrinsic excitability.

### Can we use structure to predict function *in vivo*? AIS properties and neuronal excitability

Whether on a canonical soma-origin axon or one that emanates from a dendrite, the AIS’s structural properties (distance from soma and length) can have a major impact on a neuron’s excitability. For the property of AIS position the precise nature of this impact remains unresolved, and is likely to depend on various factors including variation in neuronal morphology (Hamada et al., 2016; Parekh and Ascoli, 2015). In contrast, changes in AIS length have a much clearer corollary. Experimental and theoretical results are in close agreement that, all else being equal, a shorter AIS leads to decreased excitability (Evans et al., 2015; Goethals and Brette, 2019; Grubb and Burrone, 2010; Gulledge and Bravo, 2016; Höfflin et al., 2017; Jamann et al., 2020; Kuba et al., 2010; Pan-Vazquez et al., 2020; Sohn et al., 2019; Wefelmeyer et al., 2015). Our data showing brief sensory deprivation-induced AIS shortening and decreased excitability in OB DA neurons are entirely consistent with this coherent picture.

Importantly, while changes in both AIS position and length have been described in cultured neurons (Chand et al., 2015; Dumitrescu et al., 2016; Evans et al., 2013, 2015; Grubb and Burrone, 2010), plasticity of AIS position without any accompanying length change has yet to be described in intact networks. Indeed, all *in vivo* activity-dependent AIS plasticity described to date seems to express itself as length changes (Fig.8 here; (Höfflin et al., 2017; Jamann et al., 2020; Kuba et al., 2010; Pan-Vazquez et al., 2020)). Failure to describe *in vivo* AIS position changes could be due to a physical impediment to moving this macromolecular structure, which is tightly linked to extracellular matrix proteins (Brückner et al., 2006), when the overall 3D circuit structure is in place. Alternatively, *in vivo* AIS positional changes might be possible, but we have yet to probe the cell types that are capable of this with an appropriate manipulation. Finally, it is important to note that the main caveat of most *in vitro* and all *in vivo* AIS plasticity studies is that analysis has been done at the population level, and links between AIS and excitability changes on a cell-by-cell level are few and far between. Future studies will need to address this by pairing electrophysiological recordings with new tools for AIS live imaging (Dumitrescu et al., 2016).

### Implications for olfactory processing

We find here that 24 h sensory deprivation leaves bulbar excitatory neurons’ intrinsic excitability unchanged, but recruits structural and intrinsic plastic mechanisms in a specialized population of inhibitory interneurons, as well as producing downregulated TH levels in all DA neurons. What are the functional implications of these different neuronal responses? By releasing GABA and dopamine that can target OSN terminals, DA neurons act as gain controllers at the first synapse in olfaction (Borisovska et al., 2013; Hsia et al., 1999; Vaaga et al., 2017). Thanks to their rapid activity-dependent regulation of TH expression, both subtypes of DA cell might respond to decreased afferent input by producing and releasing less dopamine, thus decreasing feedback inhibition of OSN terminals. This could be a very effective mechanism to rapidly counterbalance the effects of sensory deprivation by increasing the gain of the first synapse in the olfactory system, potentially thereby heightening odour sensitivity. Indeed, our data represent the fastest known description of an extremely well-described phenomenon which – at least following longer-term manipulations – appears responsible for balancing bulbar input-output functions in the face of sensory deprivation (Baker et al., 1993; Cho et al., 1996; Wilson and Sullivan, 1995).

The AIS shortening and decreased excitability in axon-bearing DA cells could further accentuate the deprivation-associated relief of inhibition in the glomerular layer. Decreases in TH levels and decreases in neuronal excitability appear broadly synergistic, and together should locally increase the gain of nose-to-brain transmission. However, axon-bearing DA cells have widely arborized dendritic trees and a long-spanning axon (Banerjee et al., 2015; Galliano et al., 2018; Kiyokage et al., 2017), and are believed to contribute not only to local intraglomerular signaling and gain control, but also, by means of long-range lateral inhibition (Banerjee et al., 2015; Liu et al., 2013; Whitesell et al., 2013), to odour identification and discrimination (Linster and Cleland, 2009; Uchida et al., 2000; Urban, 2002). Decreasing their excitability might therefore be expected to produce olfactory discrimination deficits. How can we reconcile these two potentially opposing effects? One could speculate that when the network is deprived of sensory inputs, a first, fast-acting response dampening all (intra- and interglomerular) inhibition to increase overall sensitivity (Kuhlman et al., 2013) could be prioritized over maintaining fine discrimination. Then, if the sensory deprivation persists, a more nuanced solution might be implemented in which other neuron types adapt their excitability to reach a new stable network set point, whilst permitting interglomerular connections to reprise their more powerful long-range inhibitory function (Gainey and Feldman, 2017). In addition, the long-range interglomerular projections of glomerular layer DA neurons have also been proposed to underlie gain control modulation of OSN➔M/TC signaling (Banerjee et al., 2015; Bundschuh et al., 2012), so targeted decreases in their excitability could be another mechanism for ensuring maximal impact of diminished OSN inputs, especially in the initial stages once the state of deprivation begins to resolve. In this way, specific plastic changes in one cell type might shift the balance of information processing in sensory circuits to prioritise detection over discrimination when input activity is diminished.

### Homeostasis in cells or circuits? Inhibitory neurons as first responders

While not preponderant in cortex, inhibitory neurons constitute the main population in the olfactory bulb (Shepherd, 2004). Heterogeneous in all brain areas, inhibitory neurons can be just as plastic as their excitatory counterparts, but can respond differently to the same sensory input (Gainey and Feldman, 2017). Understanding this differential excitatory/inhibitory plasticity and its time course could help unpack one of the most puzzling phenomena in neuroscience: how stability and plasticity coexist to ensure both homeostasis and learning (Fox and Stryker, 2017). Indeed, one could speculate that while the plasticity of excitatory neurons is mostly Hebbian and aimed at supporting the acquisition of new associations (Bekisz et al., 2010; Gao et al., 2017; Yiu et al., 2014), one of the main functions of activity-dependent plasticity in inhibitory neurons is to act as ‘first responders’. In this scheme, plasticity in local inhibitory cells acts to compensate a short-lived change in sensory input and to maintain homeostasis – not at the single-cell level, but at the network level. If then the sensory perturbation persists and becomes the ‘new normal’, excitatory cells might need to activate homeostatic plasticity mechanisms and inhibitory neurons to downscale their own fast-acting plastic response, to reach a new network set point while maintaining an appropriate dynamic range (Gainey and Feldman, 2017; Keck et al., 2017; Turrigiano, 2012; Wefelmeyer et al., 2016). The overall circuit response to a changed sensory stimulus cannot thus be inferred by solely looking at principal neurons (Hennequin et al., 2017), or by simple arithmetic sums of plastic changes in the various neuron types, or without appreciation of the length and scope of sensory manipulation. Future studies will need to holistically address how activity-dependent plasticity is differentially expressed in inhibitory and excitatory neurons in order to shape information processing in distinct brain circuits.

## Acknowledgements

This work was supported by a Sir Henry Wellcome Fellowship (103044) to EG, a Wellcome Trust Career Development Fellowship (088301), BBSRC grant (BB/N014650/1) and ERC Consolidator Grant (725729; FUNCOPLAN) to MSG, and a Medical Research Council 4 year PhD studentship to CH. We wish to thank Mark Evans and Rosie Sammons for help with 3D tracing in Fiji, Annisa Chand for instructions on nose plug manufacture, Andres Crespo for help with olfactory epithelium samples preparation and overall excellent technical assistance. Venki Murthy and all members of the Grubb, Murthy and Galliano laboratories provided helpful discussions, while Juan Burrone and Sue Jones made invaluable comments on the manuscript.

